# Neurotransmitter receptor architecture links local neurochemistry to semantic memory

**DOI:** 10.64898/2026.10.26.762019

**Authors:** JeYoung Jung, Jeong-An Gim, Matthew A. Lambon Ralph

## Abstract

A fundamental challenge in neuroscience is understanding how the molecular and neurochemical architecture of the human cortex gives rise to sophisticated higher cognitive functions. The anterior temporal lobe (ATL) is a key hub for semantic memory, and its underlying GABAergic and glutamatergic neurochemistry supports semantic memory performance. However, the molecular receptor architecture underlying these neurochemical–cognitive relationships remains unknown. Here, we combined imaging transcriptomics and in vivo magnetic resonance spectroscopy (MRS) to investigate how neurotransmitter receptor organisation relates to local neurochemistry and semantic memory in the human ATL. We found that the ATL exhibited a distinct transcriptomically inferred receptor-subunit profile relative to control cortical regions (occipital cortex and vertex). Individual-specific spatial estimates of receptor-subunit gene expression were systematically associated with local GABA+ and Glx concentrations, revealing preferential coupling between GABAergic receptor genes and GABA+ and between glutamatergic receptor genes and Glx, alongside widespread cross-system associations between GABAergic receptor architecture and Glx. ATL GABA+ and Glx concentrations were positively associated with semantic memory performance, and GABAergic and glutamatergic receptor-subunit expression showed selective relationships with semantic memory accuracy. Crucially, mediation analyses revealed that GABA+ mediated associations between GABAergic receptor architecture and semantic performance, whereas Glx mediated both glutamatergic and widespread GABAergic receptor associations.

Together, these findings reveal a coupled molecular–neurochemical architecture of the human ATL, linking neurotransmitter receptor organisation to regional neurochemical state and individual differences in semantic memory. Our findings provide a multiscale account of how cortical molecular organisation is associated with human higher cognitive function.

## Introduction

Human cognition emerges from neural circuits whose computational properties are constrained by their molecular and neurochemical architecture. Among the most fundamental determinants of cortical computation is the interplay between glutamatergic excitation and GABAergic inhibition, which regulates neuronal gain, temporal precision, synaptic plasticity and network stability ^1–5^. Yet much of human cognitive neuroscience has focused on regional activity, connectivity and behaviour, leaving questions on how molecular receptor organisation and local neurochemical states contribute to cognitive function, largely unresolved.

Bridging these levels requires the combination of information about neurotransmitter receptor organisation with *in vivo* neurochemical measurements. Regional neurotransmitter concentrations provide a macroscopic view of local neurochemical state, whereas receptor-subunit composition constrains how those signals are translated into neurotransmission. As receptors are the biophysical gatekeepers of neurotransmission, the specific configuration of receptor and receptor-subunit families determines whether a chemical signal translates into rapid synaptic transmission, slow neuromodulation, or tonic inhibition ^6–8^. Regional receptor architecture may therefore provide an important molecular interface between the local neurochemical environment and the neural computations supporting human cognition.

Imaging transcriptomics enables spatial gene-expression data to be integrated with neuroimaging, providing a means to characterise the molecular architecture of the human cortex ^9–12^. Mapping the regional expression of neurotransmitter receptor-subunit genes can reveal distinct molecular profiles, or “receptor fingerprints,” that capture the composition and diversity of receptor systems across cortical areas ^13–15^. These molecular profiles are systematically related to macroscale cortical organisation, covarying with structural and functional properties of the human cortex ^10, 11, 16^. In particular, transcriptomic and neurotransmitter receptor profiles align with functional cortical hierarchies extending from sensorimotor regions to transmodal association cortex, suggesting that regional molecular specialisation may contribute to the functional differentiation of cortical systems supporting higher-order cognition^11, 15–17^_._

A premier example of such high-level cognition is semantic memory, the system that supports the acquisition, representation, and retrieval of conceptual knowledge ^18, 19^. Semantic memory underpins language, object recognition, reasoning, and many aspects of everyday cognition ^20–22^. Converging evidence from lesion studies, functional magnetic resonance imaging (fMRI), non-invasive brain stimulation and meta-analytic evidence identifies the ATL as the transmodal hub for this system, integrating distributed, modality-specific sensory-motor information into coherent, generalizable conceptual representations ^19, 22–32^. Importantly, ATL function depends on its local neurochemical state. In vivo MRS has shown that ATL GABA concentrations predict individual semantic performance, and subsequent work has linked ATL neurochemical profiles to task-evoked neural responses and individual sensitivity to neuromodulation ^33–36^. Together, these findings suggest that semantic memory depends not only on the macroscale organisation of the ATL but also on its local neurochemical environment. However, the molecular architecture underlying these neurochemical–cognitive relationships remain unknown.

Although *in vivo* MRS provides a unique, non-invasive approach for quantifying regional neurochemical concentrations ^37^, its signals cannot be interpreted as direct measures of synaptic neurotransmission. GABA-edited MRS yields GABA+, which comprises GABA together with co-edited macromolecular signals and is thought to reflect predominantly bulk intracellular and extrasynaptic pools rather than phasic synaptic inhibition ^38, 39^. Likewise, Glx represents the combined glutamate and glutamine signal and reflects both neurotransmission and cellular metabolism through the glutamate–glutamine cycle and associated energetic processes ^40, 41^.

Thus, MRS characterises regional neurochemical state but does not identify the molecular receptor systems associated with that state. Imaging transcriptomics provides a complementary level of analysis by estimating regional receptor-subunit gene expression, thereby enabling molecular receptor architecture to be related to in vivo GABA+ and Glx concentrations.

Here, we integrated in vivo MRS with imaging transcriptomics to investigate how transcriptomically inferred neurotransmitter receptor architecture in the ATL relates to local neurochemistry and semantic memory. Participant performed a semantic association and control, pattern matching tasks (Fig. 1A) outside of the scanner and went into the scanner for MRS on the ATL and control regions (occipital cortex (OCC) and vertex) (Fig. 1B). We first examined whether the ATL exhibits a neurotransmitter receptor-subunit expression profile distinct from control cortical regions. We next tested whether participant-specific spatial estimates of receptor-subunit gene expression were associated with MRS-measured GABA+ and Glx, with the a priori expectation of preferential coupling between GABAergic receptor genes and GABA+ and between glutamatergic receptor genes and Glx. We then examined whether GABAergic and glutamatergic receptor-subunit expression selectively relates to semantic memory performance. Finally, we tested whether regional GABA+ and Glx mediate receptor–behaviour associations. By integrating molecular, neurochemical and behavioural measures within a cortical system, this study tests a multiscale model in which local receptor architecture contributes to the neurochemical basis of human semantic memory.

**Figure 1.**
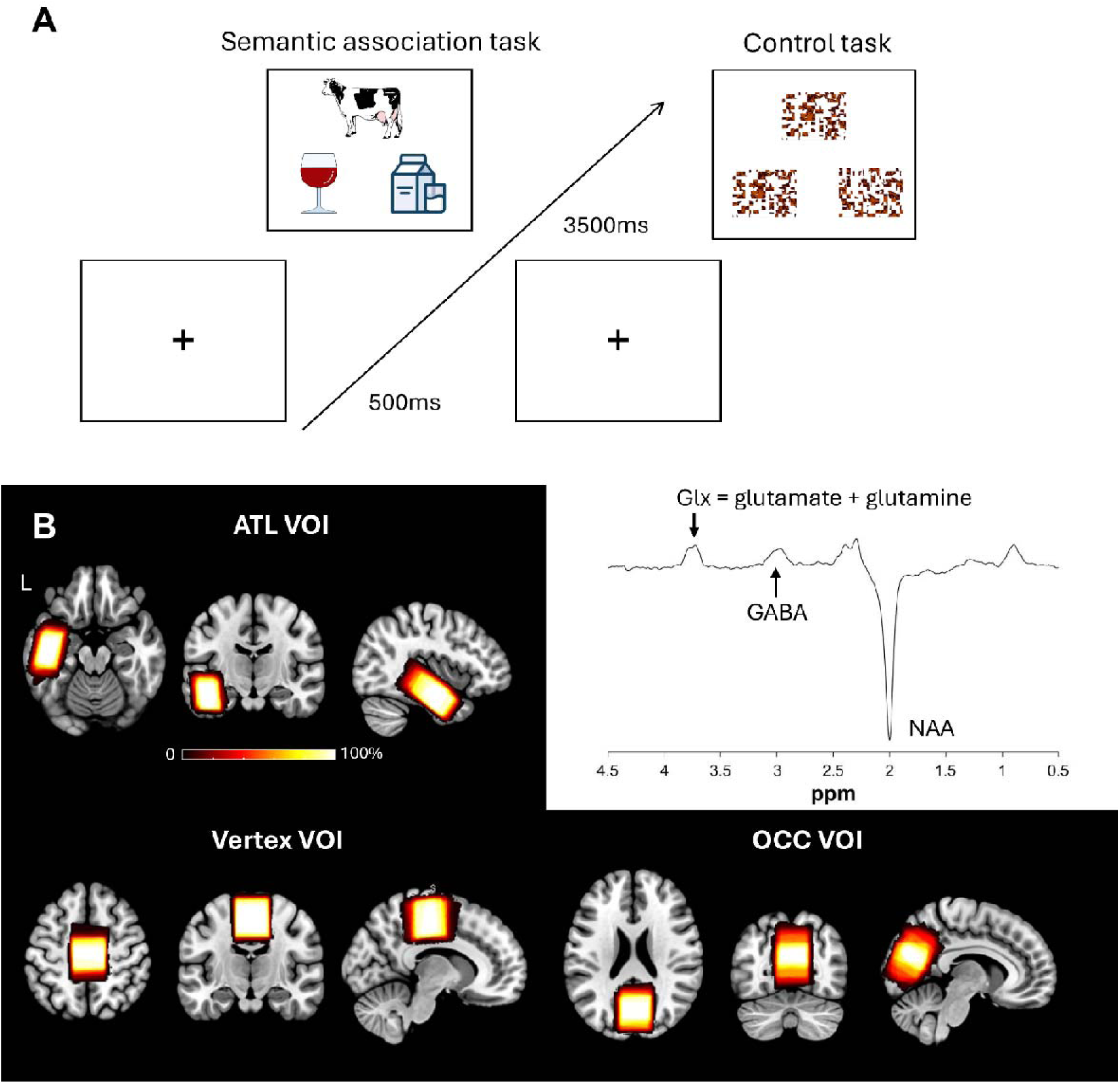
Experimental paradigm and magnetic resonance spectroscopy (MRS) volume localization. **(A)** Schematic illustration of the behavioural tasks and trial structure. Following a fixation cross (500ms), participants perform a semantic association task (left) where they match a target stimulus (e.g., a cow) to its semantically related alternative (e.g., milk) and a perceptually matched scrambled pattern matching task as a control task (right). Stimuli are presented for up to 3500ms or until a response is made. **(B)** Volume-of-interest (VOI) localization and representative MRS spectrum. Left and bottom panels display the spatial overlap maps of the planning voxels across participants for the anterior temporal lobe (ATL; top left, n = 45), vertex (control site; bottom left, n = 22), and occipital cortex (OCC; control sit; bottom right, n = 23) VOIs, overlaid on a standard MNI template. The colour bars denote the percentage of participant overlap from 0% to 100%. Right panel illustrates a representative MRS spectrum from a single participant, showing characteristic peaks for Glx (glutamate + glutamine), GABA, and NAA (N-acetylaspartate).

## Results

### The anterior temporal lobe exhibits a distinct neurotransmitter receptor architecture

To determine whether the ATL exhibits a distinct molecular receptor architecture, we used imaging transcriptomics to estimate regional neurotransmitter receptor-subunit gene expression and compared the resulting profiles with those of two control regions, the OCC and vertex (Fig. 1B). Regional neurotransmitter receptor-subunit gene expression was estimated using the InSpectro-Gadget toolbox ^42^, which interfaces with whole-brain microarray mRNA expression data from the Allen Human Brain Atlas (https://human.brain-map.org/) (Fig. 2). Participant-specific spatial masks for the ATL, vertex, and OCC in native space were normalized to MNI152 standard space. Within this standardized 3D voxel-of-interest (VOI), voxel-wise expression values were extracted for a targeted panel of receptor subunits, such as the receptor isoforms (GABRA1–4). Following robust sigmoid normalization relative to the cortical median and grey-matter filtering, the median normalized expression across valid voxels was computed for each gene to construct participant-specific transcriptomic receptor profiles on a standardized 0–1 scale.

**Figure 2.**
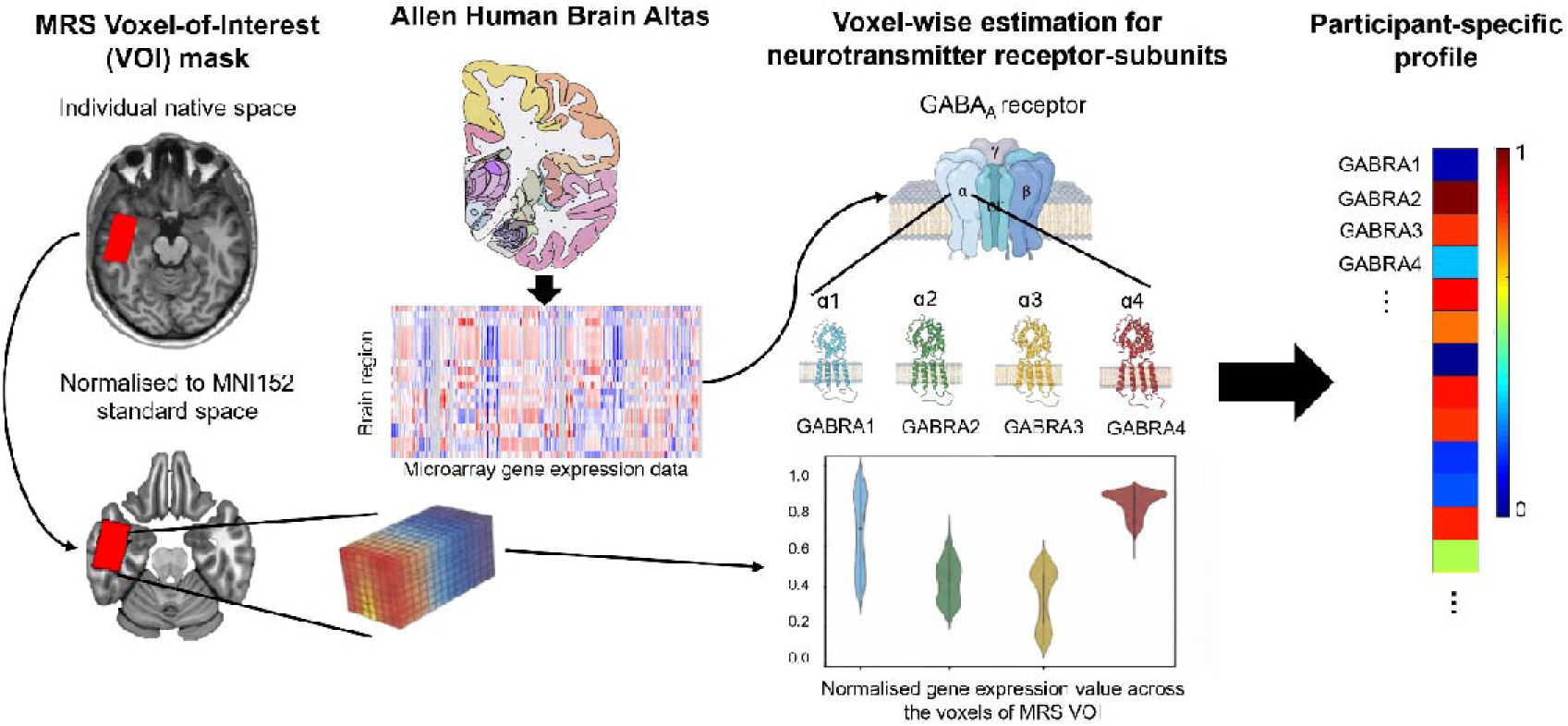
Overview of the imaging transcriptomics workflow linking magnetic resonance spectroscopy (MRS) voxel localization to participant-specific receptor profiles. The pipeline integrates macroscopic in vivo neuroimaging with microscopic transcriptomic data from the Allen Human Brain Atlas (AHBA). (Left) MRS Voxel-of-Interest (VOI) Mask: Participant-specific spatial masks (e.g., for the anterior temporal lobe) defined in individual native space are normalized to MNI152 standard space to construct a standardized 3D voxelated block. (Center-Left) Allen Human Brain Atlas & Microarray Data: Whole-brain microarray mRNA expression data are obtained from the AHBA and mapped using cortical representations in standard MNI152 space. (Center-Right) Voxel-wise Estimation for Neurotransmitter Receptor-Subunits: Voxel-level mRNA expression values are extracted for a targeted panel of receptor subunits and genes, such as GABA_A_ receptor subunits (GABRA1–4) within the VOI. Each subunit corresponds to a distinct color code matching the receptor-subunit protein models (blue for 1/GABRA1, green for 2/GABRA2, yellow for 3/GABRA3, and red/brown for 4/GABRA4), with violin plots depicting normalized gene expression values across the voxels of the MRS VOI. (Right) Participant-Specific Profile: Following robust sigmoid normalization relative to the cortical median and grey-matter filtering, median expression values across valid voxels are calculated for each gene, yielding a standardized participant-specific transcriptomic receptor profile (scaled from 0 to 1) coloured according to each respective receptor subunit for downstream statistical analyses.

Across the 75 receptor-subunit genes examined, the ATL showed a characteristic transcriptomic profile encompassing GABAergic, glutamatergic, cholinergic, adrenergic, dopaminergic and serotonergic receptor systems (Fig. 3A).

**Figure 3.**
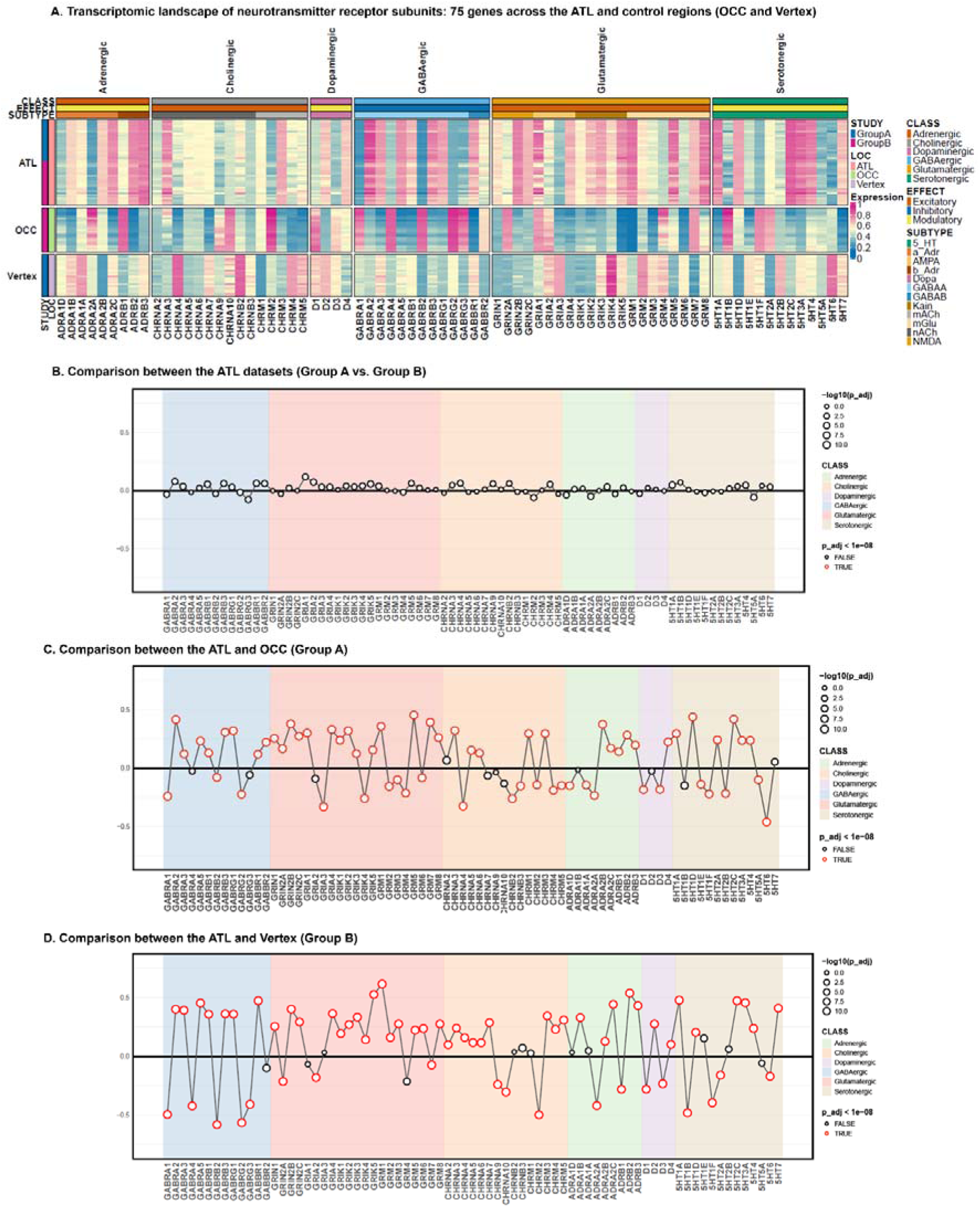
The anterior temporal lobe exhibits a distinct neurotransmitter receptor architecture. **(A)** Heatmap illustrating regional neurotransmitter receptor-subunit gene expression profiles across the anterior temporal lobe (ATL), occipital cortex (OCC), and vertex. Columns represent the 75 receptor-subunit genes grouped by neurotransmitter family and receptor subtype. Rows represent participant-specific regions of interest. Subunit genes (x-axis) are categorized by functional receptor class (glutamatergic, GABAergic, dopaminergic, serotonergic, cholinergic, adrenergic) and physiological Effect class (excitatory, inhibitory, modulatory). Metadata bars indicate dataset cohort (Group A, 23; Group B, 22) and anatomical region (ATL, OCC, & Vertex). **(B)** Differential receptor-subunit gene expression between the two ATL acquisition cohorts (Group B minus Group A). No receptor-subunit genes differ significantly. **(C)** Differential receptor-subunit gene expression between the ATL and OCC. **(D)** Differential receptor-subunit gene expression between the ATL and the vertex.

We first assessed the spatial robustness of ATL transcriptomic estimates across the two acquisition cohorts. Comparisons of median receptor-subunit gene expression between Groups A and B revealed no significant differences across any of the 75 genes (Fig. 3B). We next examined whether this receptor architecture distinguished the ATL from other cortical regions. Relative to the OCC, the ATL showed widespread differences in receptor-subunit gene expression, with a substantial proportion of the 75 genes (Fig. 3C). Differential expression encompassed multiple neurotransmitter systems, including GABAergic (*GABRA2, GABRA3, GABRB1, GABRB3,* and *GABBR1*), glutamatergic (*GRIA1, GRIA2, GRIN2A, GRIK2, GRM1, GRM5,* and *GRM7*), cholinergic (*CHRM2* and *CHRM4*), adrenergic (*ADRA2A* and *ADRB1*) and serotonergic (*HTR1A* and *HTR2C*) receptor genes. A similarly differentiated profile was observed when the ATL was compared with the vertex (Fig. 3D). Significant differences were evident across GABAergic (*GABRA2, GABRA4, GABRB1, GABRG1,* and *GABBR1*), glutamatergic (*GRIN1, GRIN2B, GRIK4, GRIK5,* and *GRM1*), adrenergic (*ADRA2A, ADRB1,* and *ADRB2*) and serotonergic (*HTR1A, HTR2A, HTR2C,* and *HTR7*) receptor systems.

Line plots display the regional difference in normalized median expression across the 75 receptor-subunit genes. Nodes represent individual genes colored by their corresponding functional class. Node size corresponds to the negative log10- transformed adjusted p-value. Red outlines indicate statistical significance surviving strict Bonferroni correction (adjusted p < 10^-8^).

### Regional receptor-subunit gene expression is associated with local neurochemical profiles

We next examined whether regional neurotransmitter receptor-subunit gene expression was associated with local neurochemical concentrations (Fig. 4 and Table S2). Partial correlation analyses controlling for age and sex revealed selective relationships between transcriptomic receptor profiles and MRS-measured GABA+ and Glx concentrations, supporting the hypothesis that molecular receptor architecture contributes to regional neurochemical variation.

**Figure 4.**
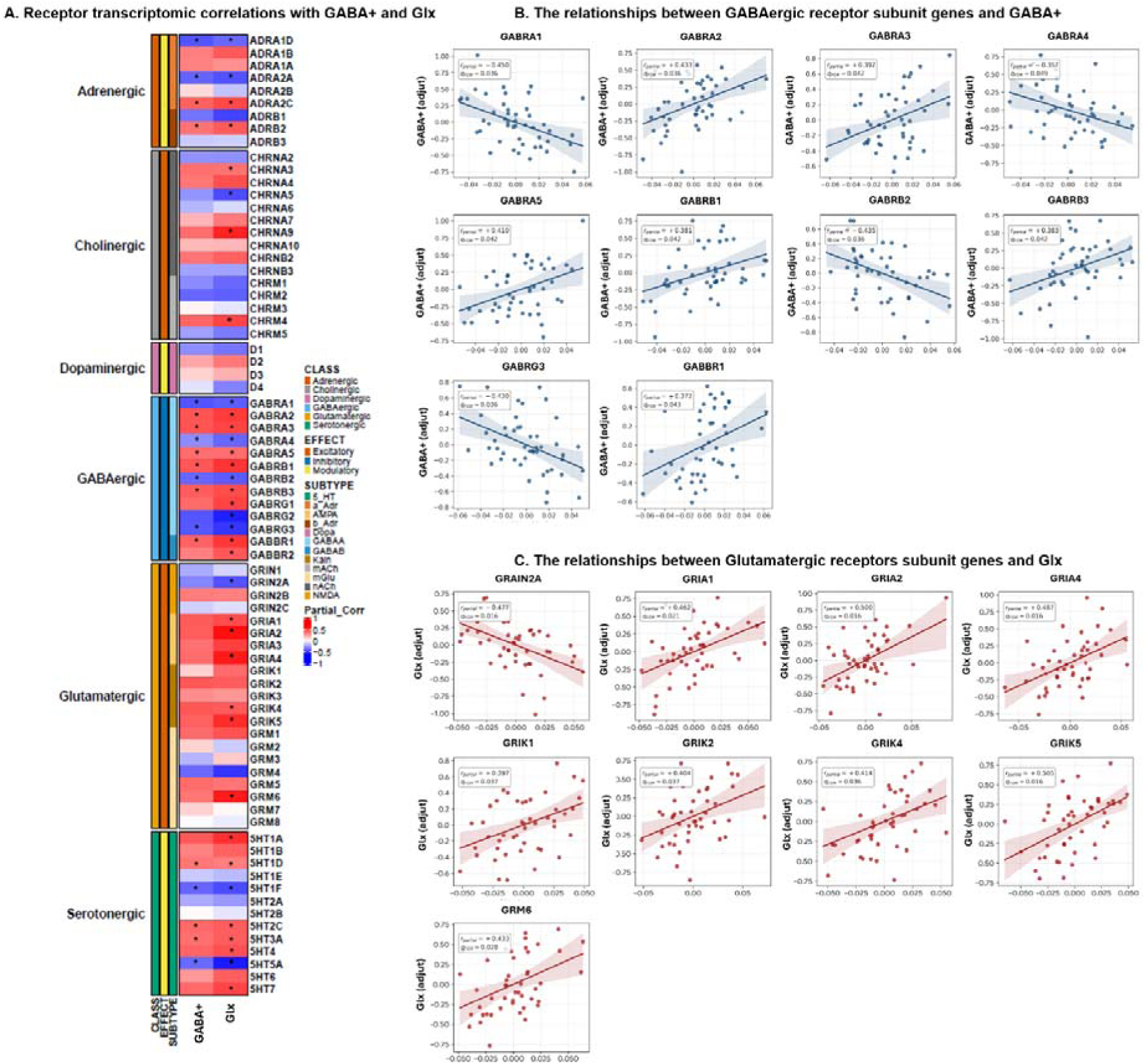
Regional neurotransmitter receptor-subunit gene expression is associated with local neurochemical profiles. **(A)** Heatmap showing partial correlation coefficients between regional neurotransmitter receptor-subunit gene expression and MRS-measured GABA+ and Glx concentrations. Partial correlations controlled for age and sex. Genes are grouped according to neurotransmitter family and receptor subtype. Colour scale indicates correlation coefficients (blue, negative correlation; red, positive correlation). Asterisks indicate significant associations following class-wise False Discovery Rate correction (* *q* _FDR_ < 0.05). **(B)** Scatter plots illustrating significant partial correlations between GABAergic subunit gene expression and GABA+ levels. **(C)** Scatter plots illustrating significant partial correlations between Glutamatergic subunit gene expression (spanning NMDA, AMPA, Kainate, and mGlu receptor families) and concentrations.

Within the GABAergic receptor family, 10 of the 13 receptor-subunit genes were significantly associated with MRS-measured GABA+ concentrations (Fig. 4B). Positive associations were observed for *GABRA2* (*r* = 0.433, *q* _FDR_ = 0.036), *GABRA3* (*r* = 0.392, *q* _FDR_ = 0.042), *GABRA5* (*r* = 0.410, *q* _FDR_ = 0.042), *GABRB1* (*r* = 0.381, *q* _FDR_ = 0.042), *GABRB3* (*r* = 0.383, *q* _FDR_ = 0.042), and *GABBR1* (*r* = 0.372, *q* _FDR_ = 0.043). In contrast, negative associations with GABA+ were observed for *GABRA1* (*r* = −0.450, *q* _FDR_ = 0.036), *GABRA4* (*r* = −0.357, *q* _FDR_ = 0.049), *GABRB2* (*r* = −0.435, *q* _FDR_ = 0.036), and *GABRG3* (*r* = −0.430, *q* _FDR_ = 0.036). Thus, regional GABA+ concentrations were related to the expression profiles of multiple GABAergic receptor subunits, although the direction of these associations differed across genes.

A corresponding relationship was observed between glutamatergic receptor architecture and Glx concentrations, with 9 glutamatergic receptor-subunit genes showing significant associations after FDR correction (Fig. 4C). Positive associations were evident for the AMPA receptor genes *GRIA1* (*r* = 0.462, *q* _FDR_ = 0.021), *GRIA2* (*r* = 0.500, *q* _FDR_ = 0.016), and *GRIA4* (*r* = 0.487, *q* _FDR_ = 0.016); the kainate receptor genes *GRIK1* (*r* = 0.397, *q* _FDR_ = 0.037), *GRIK2* (*r* = 0.404, *q* _FDR_ = 0.037), *GRIK4* (*r* = 0.414, *q* _FDR_ = 0.036), and *GRIK5* (*r* = 0.505, *q* _FDR_ = 0.016); and the metabotropic receptor gene *GRM6* (*r* = 0.433, *q* _FDR_ = 0.028). By contrast, *GRIN2A*, encoding an NMDA receptor subunit, was negatively associated with Glx (*r* = −0.477, *q* _FDR_ = 0.016).

Furthermore, widespread cross-system coupling was observed between Glx concentrations and all GABAergic subunit transcripts, with all 13 GABAergic genes showing significant associations (Fig. 4C). Among the strongest positive associations were GABRB1 (r = 0.523, *q* _FDR_ = 0.004), GABRA2 (r = 0.511, *q* _FDR_ = 0.004), and GABBR1 (r = 0.488, *q* _FDR_ = 0.007), whereas prominent negative associations were observed for GABRG3 (r = −0.529, *q* _FDR_ = 0.004), GABRA1 (r = −0.503, *q* _FDR_ = 0.005), GABRB2 (r = −0.501, *q* _FDR_ = 0.005), and GABRA4 (r = −0.471, *q* _FDR_ = 0.007).

Together, these findings reveal both neurotransmitter coupling between receptor-subunit expression and regional neurochemical pools and broader cross-system relationships between GABAergic receptor architecture and Glx, suggesting coordinated molecular–neurochemical organisation within the ATL.

Solid lines represent least-squares regression fits with shaded 95% confidence intervals.

Beyond the GABAergic and glutamatergic systems, significant associations with regional neurochemistry were also observed across several neuromodulatory receptor families. Within the cholinergic system, 4 of 15 receptor genes were associated with Glx. Positive associations were observed for *CHRNA3* (*r* = 0.444, *q* _FDR_ = 0.018), *CHRNA9* (*r* = 0.469, *q* _FDR_ = 0.014), and *CHRM4* (*r* = 0.467, *q* _FDR_ = 0.014), whereas *CHRNA5* was negatively associated with Glx (*r* = −0.565, *q* _FDR_ = 0.001).

Within the adrenergic system, 4 of 9 receptor genes were significantly associated with GABA+. *ADRA2C* (*r* = 0.416, *q* _FDR_ = 0.032) and *ADRB2* (*r* = 0.406, *q* _FDR_ = 0.032) showed positive associations, whereas *ADRA1D* (*r* = −0.412, *q* _FDR_ = 0.032) and *ADRA2A* (*r* = −0.401, *q* _FDR_ = 0.032) showed negative associations. Five adrenergic genes were also associated with Glx, with positive relationships for *ADRA2C* (*r* = 0.449, *q* _FDR_ = 0.014) and *ADRB2* (*r* = 0.493, *q* _FDR_ = 0.006), and negative relationships for *ADRA1D* (*r* = −0.475, *q* _FDR_ = 0.009), *ADRA2A* (*r* = −0.483, *q* _FDR_ = 0.009), and *ADRB1* (*r* = −0.500, *q* _FDR_ = 0.005).

The serotonergic system showed similarly widespread neurochemical associations. 5 of 13 receptor genes were associated with GABA+: *5HTR1D* (*r* = 0.422, *q* _FDR_ = 0.024), *5HTR2C* (*r* = 0.401, *q* _FDR_ = 0.035), and *5HTR3A* (*r* = 0.404, *q* _FDR_ = 0.035) showed positive associations, whereas *5HTR1F* (*r* = −0.427, *q* _FDR_ = 0.024) and *5HTR5A* (*r* = −0.417, *q* _FDR_ = 0.024) showed negative associations. 8 serotonergic receptor genes were associated with Glx, including positive associations for *5HTR1A, 5HTR1D, 5HTR2C, 5HTR3A, 5HTR4,* and *5HTR7* (all *q q* _FDR_ < 0.05) and negative associations for *5HTR5A* (*r* = −0.563, *q* _FDR_ = 0.001) and *5HTR1F* (*r* = −0.530, *q* _FDR_ = 0.002). In contrast, no dopaminergic receptor gene survived FDR correction for associations with either GABA+ or Glx (all *q* _FDR_ ≥ 0.105).

Secondary analyses of NAA, Cho, and tCr identified additional transcriptomic associations across receptor systems (Supplementary Fig. S1; Table S2; Supplementary Results). These analyses indicate that receptor-subunit expression is also related to broader aspects of regional metabolite composition and provide a specificity context for the primary GABA+ and Glx findings.

### Regional GABA+ and Glx concentrations are associated with semantic memory

We examined the relationship between ATL neurochemical concentrations and semantic performance. Partial correlation analyses controlling for age and sex revealed that semantic task accuracy was positively associated with both GABA+ and Glx concentrations (Fig. 5). Higher regional GABA+ was associated with greater semantic accuracy (r = 0.466, *q* _FDR_ = 0.024) (Fig. 5A), and higher Glx was also associated with greater semantic accuracy (r = 0.415, *q* _FDR_ = 0.032) (Fig. 5B).

**Figure 5.**
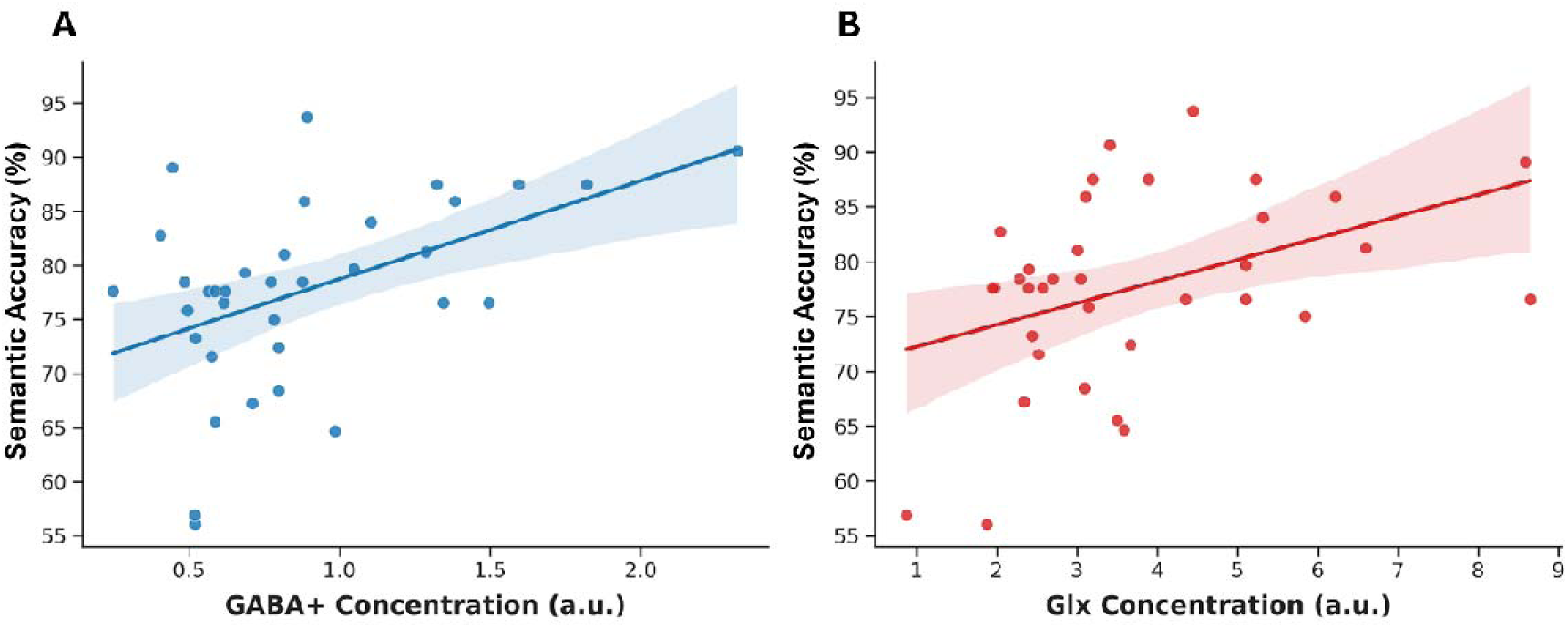
Associations between ATL neurochemistry and semantic accuracy. Partial correlations between semantic task accuracy and MRS-measured GABA+ and Glx concentrations in the ATL, controlling for age and sex. Semantic accuracy was positively associated with both GABA+ (r = 0.466, *q* _FDR_ = 0.024) and Glx (r = 0.415, *q* _FDR_ = 0.032).

Neither GABA+ nor Glx was significantly associated with semantic response time (both *q* _FDR_ > 0.05), indicating that the neurochemical relationships were specific to semantic accuracy rather than response speed. Notably, ATL GABA+ and Glx concentrations were themselves positively associated (*r* = 0.420, *p* = 0.015), indicating coordinated variation in the two regional neurochemical pools.

### Molecular receptor architecture is associated with semantic memory performance

We next examined whether regional neurotransmitter receptor-subunit gene expression was associated with behavioural performance (Fig. 6 and Table S3). Significant associations were observed selectively for semantic accuracy; no receptor-subunit genes were significant for semantic response time or for accuracy or response time on the perceptual control task. This behavioural specificity suggests that receptor–behaviour associations in the ATL were preferentially related to semantic performance rather than general task performance.

**Figure 6.**
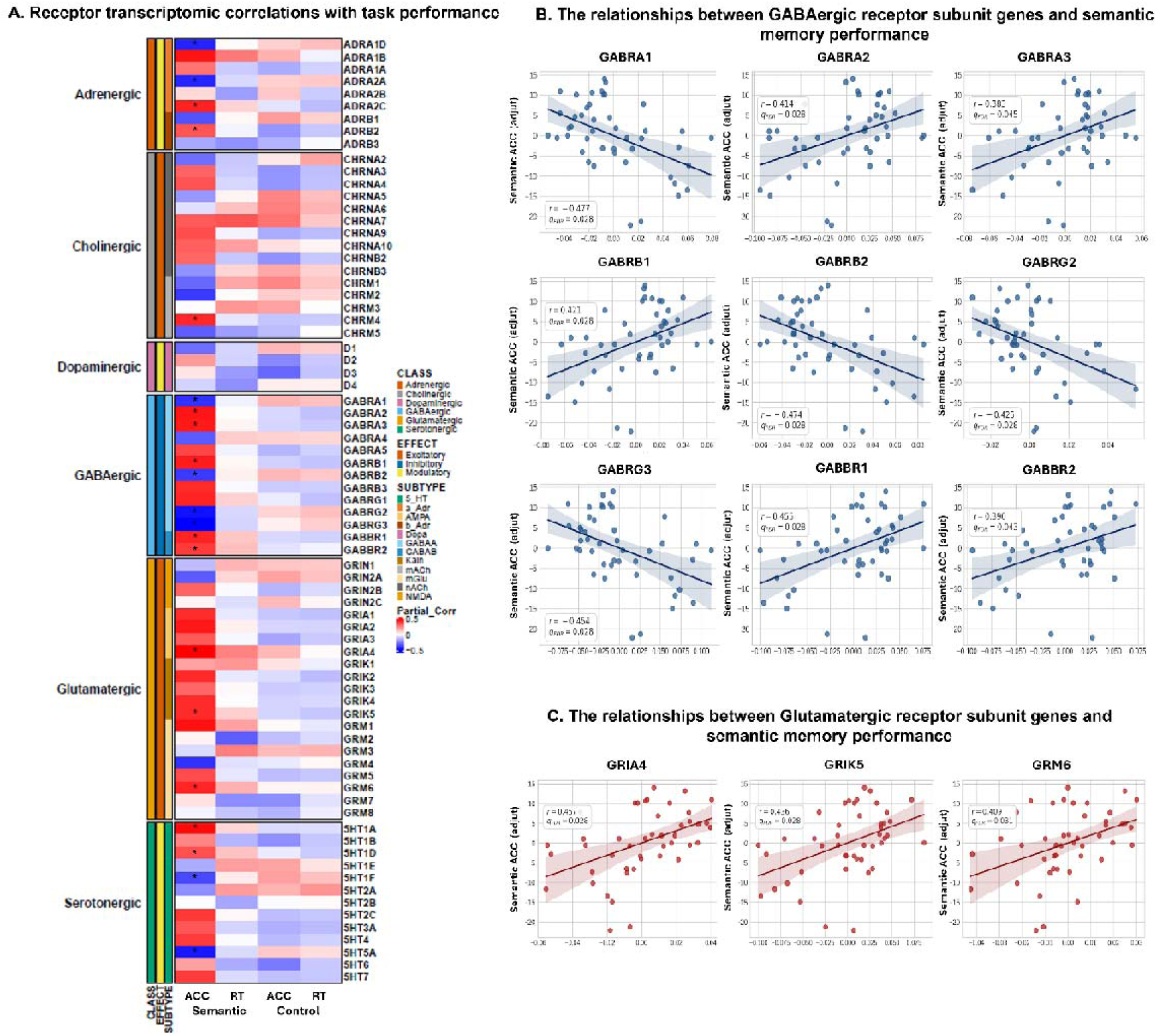
Molecular receptor architecture is associated with semantic memory performance. **(A)** Heatmap illustrating partial correlation coefficients between regional neurotransmitter receptor-subunit gene expression and task performance measures (Semantic ACC, Semantic RT, Control ACC, Control RT). Partial correlations controlled for age and sex. Receptor classes and subtypes/groups are indicated by the color-coded annotations on the left. Asterisks indicate significant associations following False Discovery Rate correction (* *q* _FDR_ < 0.05). **(B)** Scatter plots depicting the relationships between GABAergic receptor-subunit gene expression and semantic memory accuracy. **(C)** Scatter plots showing the corresponding relationships between glutamatergic receptor-subunit gene expression and semantic memory accuracy.

Within the GABAergic system, 9 receptor-subunit genes were significantly associated with semantic accuracy (Fig. 6B). Positive associations were observed for *GABRA2* (*r* = 0.414, *q* _FDR_ = 0.028), *GABRA3* (*r* = 0.383, *q* _FDR_ = 0.045), *GABRB1* (*r* = 0.421, *q* _FDR_ = 0.028), *GABBR1* (*r* = 0.455, *q* _FDR_ = 0.028), and *GABBR2* (*r* = 0.390, *q* _FDR_ = 0.043). In contrast, negative associations were observed for *GABRA1* (*r* = −0.477, *q* _FDR_ = 0.028), *GABRB2* (*r* = −0.474, *q* _FDR_ = 0.028), *GABRG2* (*r* = −0.425, *q* _FDR_ = 0.028), and *GABRG3* (*r* = −0.454, *q* _FDR_ = 0.028).

Within the glutamatergic system, 3 receptor-subunit genes were significantly and positively associated with semantic accuracy (Fig. 6C). These included the AMPA receptor gene *GRIA4* (*r* = 0.457, *q* _FDR_ = 0.028), the kainate receptor gene *GRIK5* (*r*= 0.436, *q* _FDR_ = 0.028), and the metabotropic glutamate receptor gene *GRM6* (*r* = 0.409, *q* _FDR_ = 0.031).

Solid lines represent least-squares regression fits with shaded 95% confidence intervals.

Within the adrenergic system, 4 receptor genes were significantly associated with semantic accuracy. Positive associations were observed for *ADRA2C* (*r* = 0.418, *q* _FDR_ = 0.028) and *ADRB2* (*r* = 0.403, *q* _FDR_ = 0.033), whereas *ADRA1D* (*r* = −0.465, *q* _FDR_ = 0.028) and *ADRA2A* (*r* = −0.421, *q* _FDR_ = 0.028) showed negative associations.

Within the serotonergic system, 4 receptor genes were significantly associated with semantic accuracy. Positive associations were observed for *5HTR1A* (*r* = 0.440, *q* _FDR_ = 0.028) and *5HTR1D* (*r* = 0.416, *q* _FDR_ = 0.028), whereas *5HTR1F* (*r* = −0.434, *q* _FDR_ = 0.028) and *5HTR5A* (*r* = −0.434, *q* _FDR_ = 0.028) were negatively associated with semantic accuracy.

Together, these results indicate that semantic accuracy is associated not only with GABAergic and glutamatergic receptor architecture, but also with selected adrenergic and serotonergic receptor systems, suggesting a broader neuromodulatory contribution to individual differences in semantic performance.

### Regional neurochemistry statistically mediates associations between molecular receptor architecture and semantic memory

We tested whether regional in vivo neurochemical concentrations statistically mediated associations between local neurotransmitter receptor-subunit gene expression and semantic accuracy. Mediation models included receptor-subunit gene expression as the predictor (X), MRS-measured GABA+ or Glx as the mediator (M), and semantic task accuracy as the outcome (Y), with age and sex included as covariates. The models estimated path *a* (gene expression → neurochemical mediator), path *b* (mediator → semantic accuracy controlling for gene expression), the indirect effect (*a × b*), the direct effect (*c*′), and the proportion mediated (Fig. 7A; Table S4).

**Figure 7.**
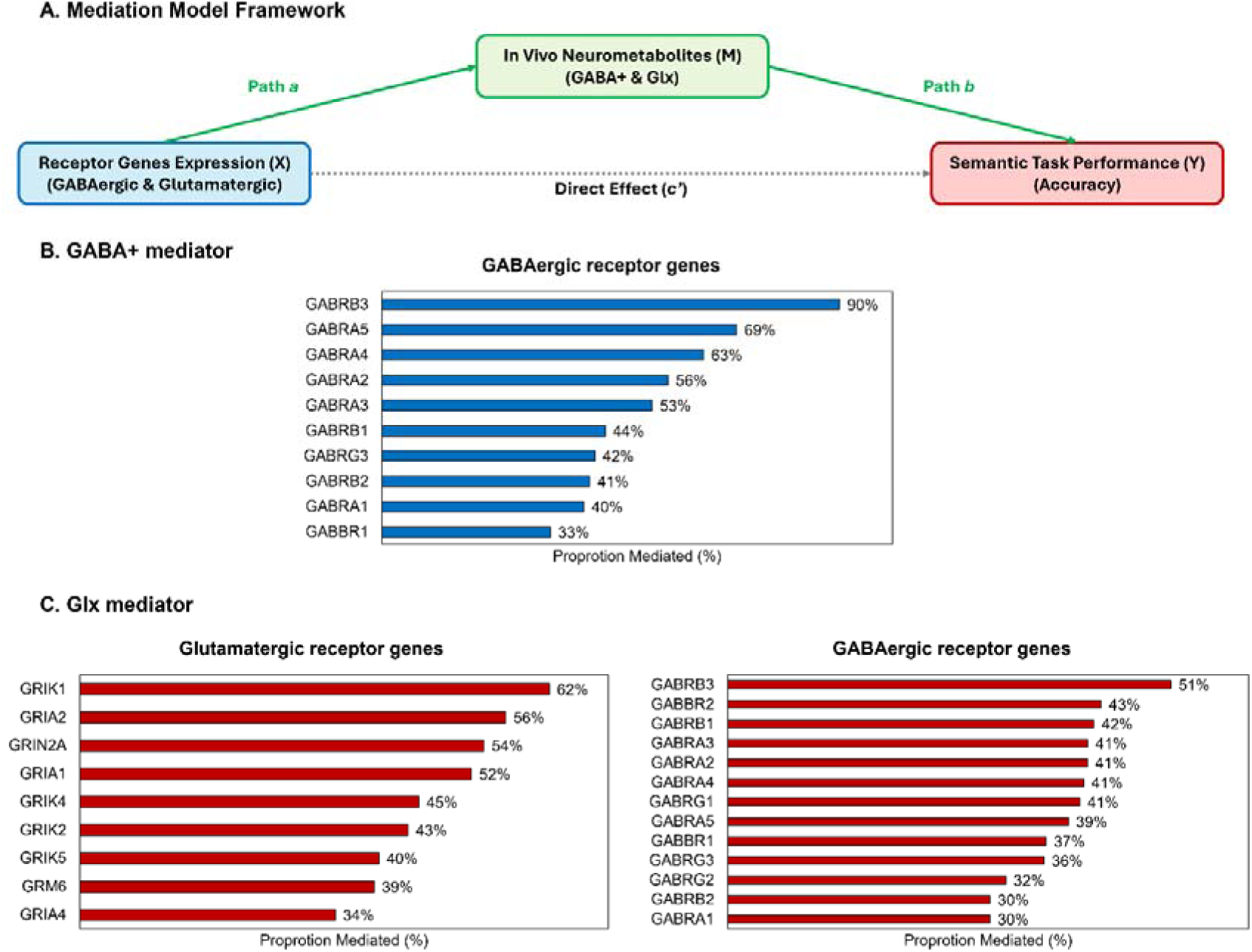
Regional neurochemical pools mediate associations between neurotransmitter receptor architecture and semantic accuracy. **(A)** Schematic of the mediation framework. Transcriptomically inferred GABAergic or glutamatergic receptor-subunit gene expression was specified as the predictor (X), MRS-measured GABA+ or Glx as the mediator (M), and semantic task accuracy as the outcome (Y). Path *a* represents the association between receptor-subunit gene expression and the neurochemical mediator, path *b* represents the association between the mediator and semantic accuracy controlling for gene expression, and *c*′ represents the direct gene–behaviour effect after accounting for the mediator. **(B)** Proportion mediated by GABA+ for GABAergic receptor-subunit genes showing significant associations with GABA+ in the preceding partial-correlation analysis. **(C)** Proportion mediated by Glx for glutamatergic (left) and GABAergic (right) receptor-subunit genes showing significant associations with Glx.

Within the GABAergic receptor system, GABA+ consistently mediated receptor– semantic associations across 10 genes (*GABRA1, GABRA2, GABRA3, GABRA4, GABRA5, GABRB1, GABRB2, GABRB3, GABRG3,* and *GABBR1*; Fig. 7B). All indirect effects were supported by 95% bootstrap confidence intervals excluding zero and accounted for an estimated 33–90% of the corresponding total gene–semantic associations. The largest proportions mediated were observed for *GABRB3* (90%), *GABRA5* (69%), *GABRA4* (63%), *GABRA2* (56%), and *GABRA3* (53%). Positive indirect effects were observed for *GABRA2, GABRA3, GABRA5, GABRB1, GABRB3,* and *GABBR1*, whereas *GABRA1, GABRA4, GABRB2,* and *GABRG3* showed negative indirect effects, consistent with the direction of their associations with GABA+. Residual direct effects (*c*′) were non-significant across these models, indicating that regional GABA+ captured a substantial component of the statistical association between GABAergic receptor architecture and semantic accuracy.

Glx also mediated receptor–semantic associations across both glutamatergic and GABAergic receptor systems (Fig. 7C). Within the glutamatergic system, 9 genes (*GRIN2A, GRIA1, GRIA2, GRIA4, GRIK1, GRIK2, GRIK4, GRIK5,* and *GRM6*) showed indirect effects with 95% bootstrap confidence intervals excluding zero. These indirect pathways accounted for an estimated 34–62% of the corresponding total associations with semantic accuracy. The largest proportions mediated were observed for *GRIK1* (62%), *GRIA2* (56%), *GRIN2A* (54%), and *GRIA1* (52%). *GRIN2A* showed a negative indirect effect, whereas the remaining glutamatergic genes showed positive indirect effects.

A widespread cross-system mediation through Glx was also observed for the GABAergic receptor system. All 13 GABAergic genes (*GABRA1–5, GABRB1–3, GABRG1–3,* and *GABBR1–2*) showed indirect effects through Glx with bootstrap confidence intervals excluding zero, accounting for an estimated 30–51% of their respective total associations with semantic accuracy. The largest proportions mediated were observed for *GABRB3* (51%), *GABRB2* (43%), *GABRB1* (42%), and *GABRA2, GABRA3, GABRA4,* and *GABRG1* (41% each).

Together, these findings demonstrated regional GABA+ and Glx as mediator linking transcriptomically inferred receptor architecture to semantic performance. GABA+ consistently mediated associations between GABAergic receptor-subunit expression and semantic accuracy, whereas Glx mediated both glutamatergic and widespread GABAergic receptor associations. This pattern supports a coupled molecular– neurochemical organisation of GABAergic and glutamatergic systems within the ATL.

## Discussion

These findings provide, to our knowledge, the first multiscale account linking neurotransmitter receptor architecture, in vivo neurochemistry and higher cognitive performance in the human cortex. By integrating imaging transcriptomics, MRS and behaviour within the ATL, we show that neurotransmitter receptor architecture is systematically related to local GABA+ and Glx concentrations, and that both molecular and neurochemical variation are associated with semantic performance. Crucially, regional neurochemistry statistically mediated neurotransmitter receptor– behaviour relationships. Thus, individual differences in semantic memory can be traced across biological scales from the molecular architecture through which neurotransmitters act, to the neurochemical milieu that shapes cortical computation, and ultimately to behaviour. Our findings suggest that this molecular–neurochemical framework moves beyond identifying the neural systems supporting semantic memory to reveal how their underlying biological organisation contributes to individual differences in human cognition.

The ATL exhibited a distinctive neurotransmitter receptor-subunit profile that differentiated it from sensory–motor regions. This extends evidence that cortical areas are distinguished not only by cytoarchitecture and connectivity, but also by systematic variation in gene expression and neurotransmitter receptor organisation ^10, 11, 13, 15, 16^. Such molecular specialisation may be particularly important in transmodal association cortex, where information must be integrated across distributed systems ^19, 43, 44^. As a central hub of the semantic network, the ATL integrates sensory, linguistic and experiential information into coherent conceptual representations ^18, 19, 45^. Our findings suggest that this functional specialisation is accompanied by a characteristic neurotransmitter receptor architecture that provides the molecular context for local cortical computation.

Critically, neurotransmitter receptor architecture was systematically related to neurochemistry measured in vivo. Most GABAergic receptor-subunit genes covaried with GABA+, whereas Glx was associated with multiple ionotropic and metabotropic glutamatergic receptor-subunit genes. These relationships bridge complementary levels of biological organisation: MRS captures regional neurochemical pools, whereas imaging transcriptomics characterises the molecular machinery through which neurotransmitters act. Recent animal and computational modelling studies further suggest that MRS-measured GABA and glutamate-related signals are linked to underlying inhibitory and excitatory neural activity ^46, 47^. Together, these findings suggest that GABA+ and Glx are embedded within the molecular organisation of the cortex, with their physiological significance shaped by the underlying neurotransmitter receptor architecture.

The GABAergic associations further reveal that bulk GABA+ does not simply scale with overall GABAergic receptor expression. GABAergic signalling spans rapid phasic inhibition, sustained tonic inhibition and slower GABA-mediated modulation ^7, 48–50^. Consistent with this diversity, some GABAergic receptor subunits were positively associated with GABA+, whereas others showed negative relationships.

Because MRS GABA+ reflects a regional neurochemical pool rather than synaptic inhibition itself ^51^, its functional significance may depend on the GABAergic neurotransmitter receptor architecture through which this neurochemical milieu acts.

A parallel organisation was evident for glutamatergic signalling. Glx covaried with AMPA, kainate, NMDA and metabotropic glutamate receptor-subunit genes, linking the regional glutamate–glutamine pool to molecular mechanisms spanning rapid excitatory transmission, synaptic plasticity and slower modulation ^3, 8, 52^. Positive associations with multiple AMPA and kainate receptor subunits suggest coordination between Glx and the molecular machinery supporting excitatory transmission, whereas the inverse relationship with *GRIN2A* highlights receptor-subtype specificity.

Together, the GABAergic and glutamatergic findings suggest that regional neurochemical pools are embedded within their underlying neurotransmitter receptor architecture, which provides the molecular context through which GABA+ and Glx modulate cortical function.

Importantly, these molecular–neurochemical relationships converged on semantic behaviour. Higher ATL GABA+ and Glx were associated with better semantic accuracy and specific GABAergic and glutamatergic neurotransmitter receptor-subunit profiles were associated with semantic performance. The GABAergic findings extend our previous observation that higher ATL GABA is associated with better semantic performance and greater selectivity of semantic-related neural responses ^33, 34^. We proposed that GABAergic inhibition sharpens distributed semantic representations by constraining weakly relevant or competing activity, thereby improving representational specificity ^1^. The present findings provide a potential molecular basis for this neurochemical–behavioural relationship. Thus, the contribution of GABA to semantic memory depends not only on the regional GABA levels but also on GABAergic neurotransmitter receptor architecture through which that neurochemical environment shapes local cortical computation.

Glutamatergic neurotransmitter receptor architecture showed a complementary relationship with semantic memory. Semantic accuracy was positively associated with *GRIA4, GRIK5* and *GRM6*, implicating both ionotropic and metabotropic glutamatergic signalling ^52, 53^. Together with the positive Glx–behaviour relationship, these findings suggest that effective excitatory transmission contributes to the activation and integration of distributed conceptual information within the ATL.

A particularly important finding was the extensive cross-system coupling between GABAergic neurotransmitter receptor architecture and Glx. GABA+ and Glx were positively related, GABAergic receptor-subunit genes showed widespread associations with Glx, and Glx statistically mediated GABAergic neurotransmitter receptor–behaviour relationships. This cross-system organisation is biologically plausible because cortical excitation and inhibition are intrinsically interdependent: excitatory activity recruits inhibitory circuits, which in turn regulate neuronal gain, timing and the propagation of excitation (Isaacson and Scanziani, 2011; Tatti et al., 2017). GABA and glutamate are also metabolically interconnected through the glutamate–glutamine–GABA cycle ^54–57^. Thus, GABAergic and glutamatergic systems may form a coupled neurotransmitter receptor–neurochemical architecture rather than operating as independent inhibitory and excitatory pathways.

Our mediation analyses bring these molecular, neurochemical and behavioural levels together. GABA+ statistically mediated relationships between GABAergic neurotransmitter receptor architecture and semantic accuracy, whereas Glx mediated both glutamatergic and widespread GABAergic receptor relationships.

Regional neurochemistry therefore emerges as a potential intermediate biological scale linking neurotransmitter receptor architecture to cognitive function. The cross-system Glx pathways further indicate that this organisation cannot be reduced to independent inhibitory and excitatory channels. Efficient semantic processing may therefore emerge from this integrated excitatory–inhibitory organisation, in which glutamatergic signalling supports the activation and integration of distributed conceptual information, while GABAergic signalling regulates its gain, selectivity and stability.

Our findings bridge a fundamental gap between molecular and cognitive neuroscience. Whereas previous human studies have related neurochemistry to neural function or behaviour, our findings show that the neurochemical milieu is itself embedded within an underlying neurotransmitter receptor architecture that is systematically related to cognitive performance. To our knowledge, this is the first study to link neurotransmitter receptor architecture, in vivo neurochemistry and behaviour within a human higher-order cognitive system. The findings therefore move beyond identifying where semantic memory occurs towards explaining how its molecular and neurochemical organisation relates to individual differences in performance. These findings provide a multiscale framework in which neurotransmitter receptor architecture provides the molecular context in which regional neurochemistry operates, bridging cortical molecular organisation with human cognition.

## Acknowledgments

The authors acknowledge Sarah Wilson, Louise Cowell, Andrew Cooper, Jan A Paul, and Mehri Kaviani for their MRI support in the project.

## Author contributions

Conceptualization: JJ; Methodology: JJ, JG; Investigation: JJ, JG; Writing— JJ, JG, MALR; Writing—review & editing: JJ, JG, MALR

## Data and materials availability

The data that support the findings of this study are available on request from the corresponding author, JJ.

## Methods and Materials

### Participants

Forty-five healthy participants (15 males, mean age = 22 ± 3 years, ranging from 19 to 33) were recruited in this study. All participants were native-English speakers with right-handed ^58^, and reported no history of neurological or psychiatric disorders. Written informed consent was obtained from all participants prior to participation. The study was approved by the ethics committee of the School of Psychology at the University of Nottingham (F1036R) and was conducted in accordance with the Declaration of Helsinki.

### Experimental design and procedure

The experimental tasks were identical to those employed in our earlier work ^33–35^. Outside the scanner, participants completed two tasks: a semantic association task and a pattern-matching task that served as a control task (Figure 1A). In the semantic association task, participants viewed a probe image displayed at the top of the screen and selected which of two images shown at the bottom was more closely related in meaning to the probe. The stimuli for this task were drawn from the *Pyramids and Palm Trees* test ^59^ and the *Camel and Cactus* test ^60^. The pattern-matching task used stimuli generated by scrambling the images from the semantic association task. Participants were required to decide which of two patterns presented at the bottom of the screen was visually identical to the probe pattern shown at the top (Figure 1A). For both tasks, responses were made by pressing one of two buttons corresponding to the available choices. Each trial began with a 500 ms fixation period, followed by stimulus presentation lasting 3500 ms. Each task consisted of 60 trials in total. PsychoPy (2024.1.6) was used to present stimuli and record responses.

### Magnetic resonance imaging acquisition

Images were collected using a General Electric (GE) SIGNA Premier 3T MR scanner equipped with a 48-channel head coil (GE Healthcare, USA). T1-weighted images were acquired using 3D MPRAGE sequence (voxel size = 1mm isotropic, field of view [FOV] = 256, matrix = 256, 256 sagittal slices, inverse time [TI] = 800ms, flip angle [FA] = 8°).

MRS data were obtained using a GABA-edited MEGA-PRESS sequence (TR =2000ms, TE = 68ms) ^37^. The voxel of interest (VOI) was manually placed in the left ventrolateral ATL (voxel size = 40 x 20 x 20mm), avoiding overlap with the hippocampus (Figure 1B). Spectra data were acquired in interleaved blocks of four averages, with the MEGA inversion pulses applied at 1.95 ppm to edit GABA signal.

Due to a scanner system upgrade during data collection, participants were acquired across two cohorts. In the first cohort (Group A: N = 23, 6 males, mean ageD=D21 D±D3.02 years, ranging from 19 to 33), 368 averages were obtained from the ATL VOI, and 224 averages were collected from a control voxel in the occipital lobe (30 × 30 × 30 mm). In the second cohort (Group B: N = 22, 9 males, mean ageD = D23α±D3.36 years, ranging from 19 to 29) 288 averages were collected from the ATL VOI, along with 112 averages from a control voxel located at the vertex (30 × 30 × 30 mm). These protocols provided reliable estimates of GABA and glx concentrations within the target region ^61, 62^. Each spectrum consisted of 4096 sample points acquired over a spectral width of 5 kHz. Differences in the number of spectral averages between cohorts reflected the scanner upgrade and subsequent optimisation of the MRS sequence for ATL acquisition. Importantly, quality-control analyses showed no significant differences in ATL spectral quality or metabolite estimates between Groups A and B, supporting the comparability of ATL measurements across acquisition protocols (Table S1).

### MRS analysis

MRS data were processed using GANNET version 3.4 ^63^. The spectra were first corrected for frequency and phase variations using spectral registration procedures and subsequently smoothed with a 3 Hz exponential line-broadening filter ^64^. The edited resonance at 3 ppm was quantified as GABA+, reflecting GABA signals that include contributions from co-edited macromolecules. This approach was adopted because macromolecular contamination cannot be fully separated from the edited GABA signal. Signals corresponding to GABA+, Glx, and unsuppressed water were modelled and quantified by integrating fitted peaks, using a single Gaussian model for GABA+ and Glx and a Gaussian-Lorentzian model for water. In addition to these metabolites, concentrations of other neurochemicals including N-acetylaspartate (NAA), choline (Cho), and total creatine (tCr) were also estimated.

Each spectroscopy voxel was coregistered to the participant’s T1-weighted anatomical image, which was segmented into grey matter (GM), white matter (WM), and cerebrospinal fluid (CSF). These segmentations were used to apply tissue composition corrections to the metabolite estimates. The correction procedure assumed negligible metabolite concentrations in CSF and a twofold higher concentration in GM relative to WM ^65^. All neurochemical concentrations were referenced to water and further adjusted using the GM/(GM + WM) ratio.

Spectral quality was evaluated on a volume-by-volume basis using several metrics, including fitting error, signal-to-noise ratio (SNR), and linewidth. Data were excluded if any metabolite exhibited a fitting error greater than 20%, if the water peak full width at half maximum exceeded 40 Hz, or if the SNR was below 20. The results of quality control and tissue segmentation are summarised in Table S1. No significant differences were observed between Group A and Group B in data quality metrics or tissue segmentation parameters, except for CSF.

### Imaging Transcriptomics

Regional neurotransmitter receptor-subunit gene expression was estimated using the open-source Python-based toolbox InSpectro-Gadget ^42^ (Python v3.14.2). This pipeline interfaces with whole-brain microarray mRNA expression data from the Allen Human Brain Atlas (https://human.brain-map.org/), utilizing MNI152 standard-space cortical representations generated via spatial interpolation of donor microarray samples ^66^.

For each participant, spatial masks for the ATL, vertex, and OCC were generated via Gannet and normalized to standard MNI152 space (Fig. 2). Within each VOI, voxel-wise mRNA expression values were extracted for a predetermined panel of 75 neurotransmitter and neuromodulator receptor-subunit genes (Table 1). Specifically, this panel encompasses 13 GABAergic genes including (L4/M_A_ *al -* 5, *βl -* 3, and *γl -* 3 subunits as well as *RI* - 2 subunits. It includes 20 glutamatergic genes covering NMDA (*GRIN1*, *GRIN2A-C*), AMPA (*GRIA1-4*), Kainate (*GRIK1-5*), and metabotropic (*GRM1-8*) receptors. The panel features 15 cholinergic genes comprising muscarinic (*CHRM1-5*) and nicotinic (*CHRNA2-7*, *CHRNA9-10*, *CHRNB2-3*) receptors. Adrenergic contributions are represented by 9 genes including *a,* (*ADRA1A-B*, *ADRA1D*), *a_2_* (*ADRA2A-C*), and *β* (*ADRB1-3*) receptor classes. Dopaminergic receptors are covered by 5 genes spanning D_1_-like *(D_1_, D_5_)* and D_2_—like *(D_2_* - 4) families. Finally, 13 serotonergic genes are included, covering *5- HT1A-F*, *5-HT2A-C*, *5-HT3A*, *5-HT4*, *5-HT5A*, *5-HT6*, and *5-HT7* receptors. Raw mRNA expression values were normalised relative to the cortical median using a robust sigmoid normalisation procedure based on the interquartile range ^67^. Voxels outside grey matter or presenting expression values below a threshold of 0.1 were treated as missing data and excluded from subsequent calculations. For each participant and region, the median normalised expression value across all valid voxels was then calculated separately for each receptor-subunit gene, yielding participant-specific spatial estimates of transcriptomic receptor architecture for subsequent statistical analyses (Fig. 2).

**Table 1.** Neurotransmitter receptor subunit genes and their functional effect classes

| Receptor Class | Subtype / Group | Effect Class & Mechanism | Targeted Genes |
| --- | --- | --- | --- |
| GABAergic | GABA<sub>A</sub> | Fast Phasic / Tonic Inhibition | *GABRA1*, *GABRA2*, *GABRA3*, *GABRA4*, *GABRA5*, *GABRB1*, *GABRB2*, *GABRB3*, *GABRG1*, *GABRG2*, *GABRG3* |
|  | GABA<sub>B</sub> | Slow Metabotropic Inhibition | *GABBR1, GABBR2* |
| Glutamatergic | NMDA | Slow/Coincident Fast Excitation | *GRIN1, GRIN2A, GRIN2B, GRIN2C* |
|  | AMPA | Fast Ionotropic Excitation | *GRIA1, GRIA2, GRIA3, GRIA4* |
|  | Kainate | Ionotropic Excitation & Presynaptic Modifying | *GRIK1, GRIK2, GRIK3, GRIK4, GRIK5* |
|  | mGlu | Metabotropic Modulatory | *GRM1, GRM2, GRM3, GRM4, GRM5, GRM6, GRM7, GRM8* |
| Cholinergic | Muscarinic (mACh) | Metabotropic Neuromodulation | *CHRM1, CHRM2, CHRM3, CHRM4, CHRM5* |
|  | Nicotinic (nACh) | Fast Ionotropic Excitation | *CHRNA2, CHRNA3, CHRNA4, CHRNA5, CHRNA6, CHRNA7, CHRNA9, CHRNA10, CHRNB2, CHRNB3* |
| Adrenergic | α-Adrenergic | Metabotropic Neuromodulation | *ADRA1A, ADRA1B, ADRA1D, ADRA2A, ADRA2B, ADRA2C* |
|  | β-Adrenergic | Metabotropic Neuromodulation | *ADRB1, ADRB2, ADRB3* |
| Dopaminergic | Dopamine | Metabotropic Neuromodulation | *D1, D2, D3, D4, D5* |
| Serotonergic | 5-HT | Mixed Neuromodulation | *HTR1A, HTR1B, HTR1D, HTR1E, HTR1F, HTR2A, HTR2B, HTR2C, HTR3A, HTR4, HTR5A, HTR6, HTR7* |

### Statistical analysis

Regional transcriptomic receptor profiles were compared between the ATL and each control region (OCC and vertex) using *t*-tests. ATL receptor-subunit expression estimates were also compared between the two acquisition cohorts to assess the spatial robustness of transcriptomic estimates across independently acquired voxel distributions. Multiple comparisons across the 75 receptor-subunit genes were controlled using Bonferroni correction, corresponding to a significance threshold of *p* < 6.67 × 10-

Associations between participant-specific spatial estimates of receptor-subunit gene expression and MRS-measured neurochemical concentrations (GABA+, Glx, N-acetylaspartate [NAA], choline [Cho], and total creatine [tCr]) were assessed using partial correlations controlling for age and sex. GABA+ and Glx constituted the primary neurochemical measures. For these analyses, Benjamini-Hochberg false discovery rate (FDR) correction was applied separately within each prespecified neurotransmitter receptor family (GABAergic, glutamatergic, cholinergic, adrenergic, dopaminergic, and serotonergic), with FDR-adjusted *q* < 0.05 considered significant. Secondary analyses of NAA, Cho, and tCr were corrected across the full panel of 75 receptor-subunit genes.

To establish the relationship between regional neurochemistry and semantic performance, partial correlations controlling for age and sex were performed between ATL GABA+ and Glx concentrations and semantic task performance), with FDR-adjusted *q* < 0.05 considered significant. The association between GABA+ and Glx concentrations was examined using the same approach.

Associations between receptor-subunit gene expression and behavioural performance were assessed using partial correlations controlling for age and sex. Analyses were conducted separately for the semantic association and perceptual control tasks. Multiple comparisons were controlled using Benjamini-Hochberg FDR correction separately within each prespecified neurotransmitter receptor family, with FDR-adjusted *q* < 0.05 considered significant.

Finally, to test whether regional neurochemical pools statistically mediated associations between molecular receptor architecture and semantic memory, simple mediation analyses were performed using the PROCESS macro for SPSS (Model 4). Candidate receptor-subunit genes were selected from the preceding partial-correlation analyses based on significant FDR-corrected associations with the corresponding MRS-measured neurochemical pool. A significant gene-behaviour association was not required for inclusion in the mediation analysis. For each model, receptor-subunit gene expression was entered as the predictor (X), MRS-measured GABA+ or Glx as the mediator (M), and semantic task accuracy as the outcome (Y), with age and sex included as covariates in both the mediator and outcome models. The mediation framework estimated path *a*, representing the association between receptor-subunit gene expression and the neurochemical mediator; path *b*, representing the association between the mediator and semantic accuracy while controlling for gene expression; the total effect (*c*) of gene expression on semantic accuracy; and the direct effect (*c*) after accounting for the mediator. The indirect effect was quantified as the product of paths *a* and *b* (*a × b*). Indirect effects were estimated using 5,000 bootstrap resamples with 95% bootstrap confidence intervals (CIs) and were considered statistically supported when the 95% CI did not include zero. Because an indirect effect can occur in the absence of a significant total effect, significance of the *c* path was not required for inference regarding mediation.

For descriptive purposes, the relative magnitude of the indirect pathway was additionally expressed as the proportion mediated (PM), calculated as

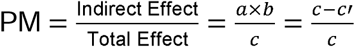

where *a* is the regression coefficient for *X -÷ M, b* is the regression coefficient for *X — Y* (controlling for X), *c* is the total effect *(Y — Y* without M), and *c’* is the direct effect *(X ^Y* controlling for M). Because PM can be unstable when the total effect is small or when direct and indirect effects differ in direction, statistical inference regarding mediation was based primarily on the bootstrapped indirect effect and its 95% CI.

