## Supplementary Information for "Neurotransmitter receptor architecture links local neurochemistry to semantic memory"

Supplementary Results

Supplementary Figure 1

Supplementary Table 1

Supplementary Table 2

Supplementary Table 3

Supplementary Table 4

**Supplementary Results**

Partial correlations between ATL transcriptomic profiles and other neurometabolites ($\text{Cr}$, $\text{Cho}$, and $\text{NAA}$) revealed dense transcriptomic associations (Figure S1 and Table S2). Cr significantly correlated with 55 of the 75 candidate genes (73.3%), spanning 12 GABAergic genes, 15 glutamatergic genes, 10 cholinergic genes, 5 adrenergic genes, 2 dopaminergic genes (*DRD1*: $r=-0.393,q_{\text{FDR}}=0.0300$; *DRD4*: $r=-0.177,q_{\text{FDR}}=0.0004$), and 11 serotonergic genes. Peak associations with $\text{Cr}$ were observed for *GABRG3* ($r=-0.581,q_{\text{FDR}}=0.0375$), *GABRA1* ($r=-0.574,q_{\text{FDR}}=0.0011$), *ADRA1D* ($r=-0.567,q_{\text{FDR}}=0.0003$), *GRIK5* ($r=0.553,q_{\text{FDR}}=0.0003$), and *GABBR1* ($r=0.551,q_{\text{FDR}}=0.0002$).

Cho significantly associated with 49 of 75 genes (65.3%), including 12 GABAergic, 15 glutamatergic, 8 cholinergic, 5 adrenergic, 1 dopaminergic (*DRD4*: $r=-0.156,q_{\text{FDR}}=0.0034$), and 8 serotonergic genes. Highest magnitude correlations with $\text{Cho}$ occurred for *GABRA2* ($r=0.655,q_{\text{FDR}}=0.0066$), *GABRB1* ($r=0.650,q_{\text{FDR}}=0.0120$), *GABRG3* ($r=-0.642,q_{\text{FDR}}=0.0182$), *GABRB2* ($r=-0.637,q_{\text{FDR}}=0.0030$), *5HT1F* ($r=-0.633,q_{\text{FDR}}=0.0006$), *ADRB2* ($r=0.633,q_{\text{FDR}}=0.5660$), *GABRA1* ($r=-0.631,q_{\text{FDR}}=0.0067$), and *GRIA2* ($r=0.623,q_{\text{FDR}}=0.0314$).

NAA, a marker of neuronal integrity and mitochondrial function, correlated significantly with 45 of 75 genes (60.0%), comprising 12 GABAergic, 12 glutamatergic, 7 cholinergic, 5 adrenergic, 1 dopaminergic (*DRD1*: $r=-0.383,q_{\text{FDR}}=0.0321$), and 8 serotonergic genes. Strongest positive correlations with $\text{NAA}$ were observed for *GABBR1* ($r=0.537,q_{\text{FDR}}=0.0292$), *5HT1D* ($r=0.531,q_{\text{FDR}}=0.0062$), *ADRA2C* ($r=0.523,q_{\text{FDR}}=0.0062$), *GABBR2* ($r=0.518,q_{\text{FDR}}=0.0062$), *GABRB1* ($r=0.513,q_{\text{FDR}}=0.0652$), and *GABRA2* ($r=0.510,q_{\text{FDR}}=0.0062$), whereas *GABRA1* ($r=-0.556,q_{\text{FDR}}=0.0062$), *5HT5A* ($r=-0.546,q_{\text{FDR}}=0.0062$), *ADRA2A* ($r=-0.535,q_{\text{FDR}}=0.0062$), and *GABRG3* ($r=-0.534,q_{\text{FDR}}=0.0233$) exhibited strong negative correlations.

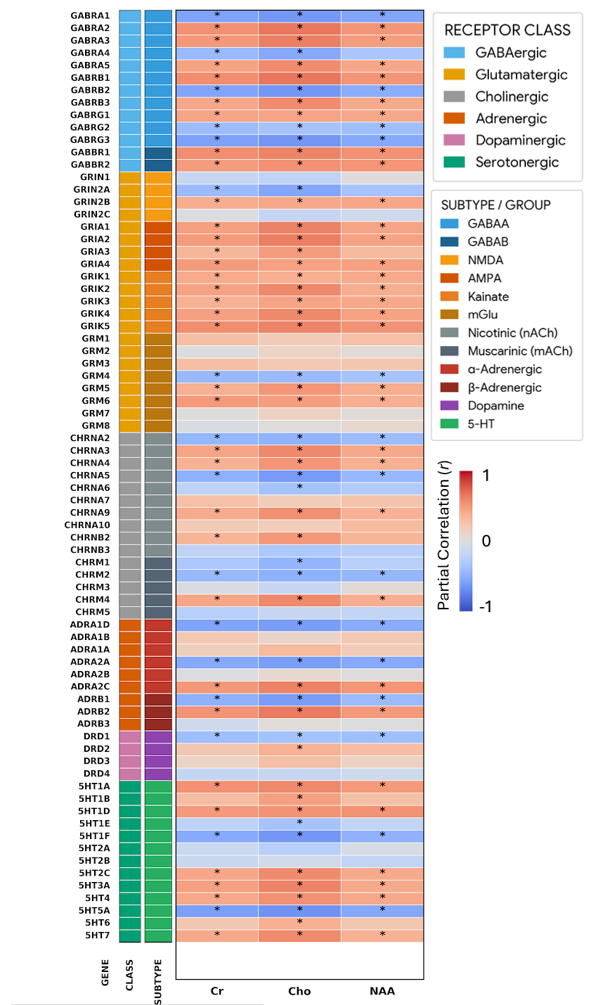
**Supplementary Figure 1**

Figure S1. Regional neurotransmitter receptor-subunit gene expression is associated with additional neurometabolite profiles.

**Supplementary Table 1**

| Region | N | Data quality metrics | | | | | Tissue segmentations | | |
| --- | --- | --- | --- | --- | --- | --- | --- | --- | --- |
|  |  | Water | | | GABA+ | Glx |  |  |  |
|  |  | Linewidth (FWHM) | SNR | Fit error (%) | Fit error (%) | Fit error (%) | GM (%) | WM (%) | CSF (%) |
| ATL | 38 | 26.13 | 23291.50 | 0.85 | 8.05 | 6.83 | 0.49 | 0.49 | 0.02 |
|  |  | (7.31) | (6446.43) | (0.52) | (2.99) | (3.09) | (0.03) | (0.04) | (0.02) |
| Group A | 18 | 28.62 | 21927.79 | 0.96 | 9.08 | 7.54 | 0.50 | 0.49 | 0.01 |
|  |  | (7.23) | (6270.88) | (0.53) | (3.59) | (1.52) | (0.04) | (0.04) | (0.01) |
| Group B | 20 | 23.40 | 24791.57 | 0.73 | 7.17 | 6.23 | 0.49 | 0.48 | 0.03 |
|  |  | (6.53) | (6456.71) | (0.50) | (2.08) | (3.91) | (0.03) | (0.04) | (0.02) |
| Vertex | 22 | 9.78 | 105151.88 | 0.49 | 5.04 | 4.60 | 0.55 | 0.35 | 0.10 |
|  |  | (2.07) | (24889.34) | (0.11) | (1.45) | (1.55) | (0.05) | (0.07) | (0.04) |
| OCC | 21 | 10.72 | 83538.72 | 0.45 | 2.63 | 2.35 | 0.70 | 0.23 | 0.08 |
|  |  | (0.83) | (25505.20) | (0.05) | (0.63) | (0.59) | (0.03) | (0.03) | (0.03) |

Table S1. The summary of MRS data quality metrics and tissue segmentation information. Data are shown as the mean (SD).

**Supplementary Table 2**

| Receptor Class | Subtype/Group |  | GABA | | Glx | | Cr | | Cho | | NAA | |
| --- | --- | --- | --- | --- | --- | --- | --- | --- | --- | --- | --- | --- |
|  |  |  | r | qFDR | r | qFDR | r | qFDR | r | qFDR | r | qFDR |
| GABAergic | GABA_A_ | GABRA1 | -0.450 | 0.036 | -0.503 | 0.005 | -0.574 | 0.004 | -0.631 | 0.002 | -0.556 | 0.005 |
|  |  | GABRA2 | 0.433 | 0.036 | 0.511 | 0.004 | 0.543 | 0.004 | 0.655 | 0.002 | 0.510 | 0.005 |
|  |  | GABRA3 | 0.392 | 0.042 | 0.446 | 0.010 | 0.505 | 0.004 | 0.618 | 0.002 | 0.482 | 0.009 |
|  |  | GABRA4 | -0.357 | 0.049 | -0.471 | 0.007 | -0.428 | 0.015 | -0.538 | 0.002 | -0.332 | 0.066 |
|  |  | GABRA5 | 0.410 | 0.042 | 0.412 | 0.013 | 0.448 | 0.011 | 0.568 | 0.002 | 0.426 | 0.023 |
|  |  | GABRB1 | 0.381 | 0.042 | 0.523 | 0.004 | 0.552 | 0.004 | 0.650 | 0.002 | 0.513 | 0.005 |
|  |  | GABRB2 | -0.435 | 0.036 | -0.501 | 0.005 | -0.556 | 0.004 | -0.637 | 0.002 | -0.514 | 0.005 |
|  |  | GABRB3 | 0.383 | 0.042 | 0.413 | 0.013 | 0.423 | 0.015 | 0.566 | 0.002 | 0.410 | 0.023 |
|  |  | GABRG1 | 0.256 | 0.155 | 0.356 | 0.033 | 0.438 | 0.013 | 0.443 | 0.008 | 0.419 | 0.023 |
|  |  | GABRG2 | -0.240 | 0.172 | -0.413 | 0.013 | -0.409 | 0.019 | -0.353 | 0.037 | -0.394 | 0.029 |
|  |  | GABRG3 | -0.430 | 0.036 | -0.529 | 0.004 | -0.581 | 0.004 | -0.642 | 0.002 | -0.534 | 0.005 |
|  | GABA_B_ | GABBR1 | 0.372 | 0.043 | 0.488 | 0.007 | 0.551 | 0.004 | 0.601 | 0.002 | 0.537 | 0.005 |
|  |  | GABBR2 | 0.331 | 0.066 | 0.415 | 0.013 | 0.482 | 0.007 | 0.533 | 0.002 | 0.518 | 0.005 |
| Glutamatergic | NMDA | GRIN1 | -0.157 | 0.524 | -0.128 | 0.565 | -0.153 | 0.406 | -0.210 | 0.246 | 0.013 | 0.960 |
|  |  | GRIN2A | -0.327 | 0.168 | -0.477 | 0.016 | -0.420 | 0.016 | -0.549 | 0.002 | -0.345 | 0.054 |
|  |  | GRIN2B | 0.292 | 0.195 | 0.248 | 0.203 | 0.392 | 0.024 | 0.415 | 0.014 | 0.424 | 0.023 |
|  |  | GRIN2C | -0.133 | 0.578 | 0.016 | 0.959 | -0.022 | 0.931 | -0.167 | 0.357 | -0.104 | 0.598 |
|  | AMPA | GRIA1 | 0.403 | 0.126 | 0.462 | 0.021 | 0.463 | 0.009 | 0.602 | 0.002 | 0.438 | 0.020 |
|  |  | GRIA2 | 0.406 | 0.126 | 0.500 | 0.016 | 0.492 | 0.004 | 0.623 | 0.002 | 0.451 | 0.017 |
|  |  | GRIA3 | 0.297 | 0.195 | 0.298 | 0.126 | 0.343 | 0.049 | 0.483 | 0.004 | 0.320 | 0.077 |
|  |  | GRIA4 | 0.321 | 0.168 | 0.487 | 0.016 | 0.491 | 0.004 | 0.455 | 0.007 | 0.458 | 0.016 |
|  | Kainate | GRIK1 | 0.179 | 0.465 | 0.397 | 0.037 | 0.404 | 0.020 | 0.385 | 0.023 | 0.408 | 0.024 |
|  |  | GRIK2 | 0.384 | 0.131 | 0.404 | 0.037 | 0.450 | 0.011 | 0.575 | 0.002 | 0.394 | 0.029 |
|  |  | GRIK3 | 0.285 | 0.195 | 0.258 | 0.194 | 0.372 | 0.032 | 0.442 | 0.008 | 0.418 | 0.023 |
|  |  | GRIK4 | 0.420 | 0.126 | 0.414 | 0.036 | 0.463 | 0.009 | 0.576 | 0.002 | 0.447 | 0.017 |
|  |  | GRIK5 | 0.358 | 0.140 | 0.505 | 0.016 | 0.553 | 0.004 | 0.600 | 0.002 | 0.527 | 0.005 |
|  | mGlu | GRM1 | 0.107 | 0.602 | 0.177 | 0.396 | 0.284 | 0.113 | 0.195 | 0.284 | 0.278 | 0.126 |
|  |  | GRM2 | 0.129 | 0.578 | -0.044 | 0.885 | -0.014 | 0.952 | 0.121 | 0.496 | 0.036 | 0.879 |
|  |  | GRM3 | 0.033 | 0.895 | 0.318 | 0.102 | 0.287 | 0.110 | 0.215 | 0.236 | 0.234 | 0.202 |
|  |  | GRM4 | -0.207 | 0.386 | -0.360 | 0.065 | -0.434 | 0.014 | -0.428 | 0.011 | -0.354 | 0.048 |
|  |  | GRM5 | 0.353 | 0.140 | 0.346 | 0.074 | 0.375 | 0.032 | 0.515 | 0.002 | 0.391 | 0.029 |
|  |  | GRM6 | 0.238 | 0.306 | 0.433 | 0.028 | 0.496 | 0.004 | 0.471 | 0.004 | 0.387 | 0.032 |
|  |  | GRM7 | 0.120 | 0.582 | -0.009 | 0.959 | 0.000 | 1.000 | 0.135 | 0.449 | 0.025 | 0.919 |
|  |  | GRM8 | 0.023 | 0.897 | -0.094 | 0.681 | -0.033 | 0.888 | 0.003 | 0.987 | 0.073 | 0.731 |
| Cholinergic | Nicotinic (nACh) | CHRNA2 | -0.247 | 0.265 | -0.298 | 0.128 | -0.447 | 0.011 | -0.476 | 0.004 | -0.411 | 0.023 |
|  |  | CHRNA3 | 0.404 | 0.141 | 0.444 | 0.026 | 0.438 | 0.013 | 0.577 | 0.002 | 0.416 | 0.023 |
|  |  | CHRNA4 | 0.340 | 0.141 | 0.361 | 0.066 | 0.372 | 0.032 | 0.522 | 0.002 | 0.382 | 0.032 |
|  |  | CHRNA5 | -0.326 | 0.141 | -0.565 | 0.015 | -0.486 | 0.007 | -0.613 | 0.002 | -0.445 | 0.017 |
|  |  | CHRNA6 | -0.229 | 0.288 | -0.175 | 0.355 | -0.211 | 0.246 | -0.367 | 0.030 | -0.292 | 0.110 |
|  |  | CHRNA7 | 0.130 | 0.580 | 0.241 | 0.236 | 0.276 | 0.122 | 0.187 | 0.304 | 0.254 | 0.163 |
|  |  | CHRNA9 | 0.334 | 0.141 | 0.469 | 0.020 | 0.401 | 0.020 | 0.546 | 0.002 | 0.366 | 0.041 |
|  |  | CHRNA10 | 0.111 | 0.615 | 0.209 | 0.276 | 0.205 | 0.255 | 0.180 | 0.319 | 0.316 | 0.081 |
|  |  | CHRNB2 | 0.339 | 0.141 | 0.368 | 0.066 | 0.348 | 0.047 | 0.503 | 0.002 | 0.350 | 0.051 |
|  |  | CHRNB3 | -0.201 | 0.348 | -0.212 | 0.276 | -0.205 | 0.255 | -0.320 | 0.063 | -0.262 | 0.151 |
|  | Muscarinic (mACh) | CHRM1 | -0.300 | 0.159 | -0.384 | 0.063 | -0.317 | 0.075 | -0.454 | 0.007 | -0.257 | 0.159 |
|  |  | CHRM2 | -0.318 | 0.141 | -0.351 | 0.068 | -0.432 | 0.014 | -0.485 | 0.004 | -0.453 | 0.016 |
|  |  | CHRM3 | -0.047 | 0.791 | -0.139 | 0.420 | -0.037 | 0.884 | -0.152 | 0.397 | 0.041 | 0.864 |
|  |  | CHRM4 | 0.384 | 0.141 | 0.467 | 0.020 | 0.436 | 0.014 | 0.575 | 0.002 | 0.391 | 0.029 |
|  |  | CHRM5 | -0.086 | 0.673 | -0.152 | 0.403 | -0.256 | 0.158 | -0.151 | 0.397 | -0.192 | 0.296 |
| Adrenergic | ɑ-Adrenergic | ADRA1D | -0.412 | 0.043 | -0.475 | 0.007 | -0.567 | 0.004 | -0.606 | 0.002 | -0.518 | 0.005 |
|  |  | ADRA1B | 0.094 | 0.673 | 0.135 | 0.558 | 0.218 | 0.237 | 0.108 | 0.542 | 0.214 | 0.246 |
|  |  | ADRA1A | 0.242 | 0.252 | 0.197 | 0.374 | 0.162 | 0.382 | 0.315 | 0.067 | 0.192 | 0.296 |
|  |  | ADRA2A | -0.401 | 0.043 | -0.483 | 0.007 | -0.538 | 0.004 | -0.598 | 0.002 | -0.535 | 0.005 |
|  |  | ADRA2B | 0.211 | 0.297 | -0.017 | 0.928 | -0.013 | 0.952 | 0.077 | 0.660 | 0.007 | 0.967 |
|  |  | ADRA2C | 0.416 | 0.043 | 0.449 | 0.011 | 0.503 | 0.004 | 0.607 | 0.002 | 0.523 | 0.005 |
|  | β-Adrenergic | ADRB1 | -0.369 | 0.058 | -0.500 | 0.007 | -0.487 | 0.007 | -0.606 | 0.002 | -0.420 | 0.023 |
|  |  | ADRB2 | 0.406 | 0.043 | 0.493 | 0.007 | 0.527 | 0.004 | 0.633 | 0.002 | 0.499 | 0.005 |
|  |  | ADRB3 | 0.005 | 0.979 | 0.016 | 0.928 | -0.106 | 0.567 | 0.021 | 0.913 | -0.011 | 0.960 |
| Dopaminergic | Dopamine | DRD1 | -0.213 | 0.360 | -0.325 | 0.142 | -0.393 | 0.023 | -0.365 | 0.031 | -0.383 | 0.032 |
|  |  | DRD2 | 0.249 | 0.360 | 0.255 | 0.177 | 0.222 | 0.230 | 0.370 | 0.029 | 0.292 | 0.110 |
|  |  | DRD3 | 0.195 | 0.360 | 0.209 | 0.222 | 0.124 | 0.497 | 0.282 | 0.106 | 0.136 | 0.478 |
|  |  | DRD4 | -0.021 | 0.905 | -0.305 | 0.142 | -0.177 | 0.336 | -0.156 | 0.390 | -0.107 | 0.593 |
| Serotonergic | 5-HT | 5HT1A | 0.332 | 0.089 | 0.467 | 0.010 | 0.535 | 0.004 | 0.574 | 0.002 | 0.499 | 0.005 |
|  |  | 5HT1B | 0.299 | 0.124 | 0.352 | 0.051 | 0.299 | 0.096 | 0.461 | 0.005 | 0.289 | 0.110 |
|  |  | 5HT1D | 0.422 | 0.049 | 0.465 | 0.010 | 0.497 | 0.004 | 0.530 | 0.002 | 0.531 | 0.005 |
|  |  | 5HT1E | -0.255 | 0.173 | -0.260 | 0.148 | -0.211 | 0.246 | -0.359 | 0.035 | -0.188 | 0.301 |
|  |  | 5HT1F | -0.427 | 0.049 | -0.530 | 0.007 | -0.528 | 0.004 | -0.633 | 0.002 | -0.477 | 0.009 |
|  |  | 5HT2A | -0.140 | 0.465 | -0.196 | 0.273 | -0.129 | 0.485 | -0.251 | 0.158 | -0.054 | 0.811 |
|  |  | 5HT2B | -0.018 | 0.922 | -0.084 | 0.624 | -0.135 | 0.469 | -0.087 | 0.625 | -0.193 | 0.296 |
|  |  | 5HT2C | 0.401 | 0.049 | 0.425 | 0.019 | 0.427 | 0.015 | 0.563 | 0.002 | 0.410 | 0.023 |
|  |  | 5HT3A | 0.404 | 0.049 | 0.487 | 0.010 | 0.462 | 0.009 | 0.595 | 0.002 | 0.407 | 0.024 |
|  |  | 5HT4 | 0.366 | 0.069 | 0.366 | 0.046 | 0.415 | 0.017 | 0.539 | 0.002 | 0.421 | 0.023 |
|  |  | 5HT5A | -0.417 | 0.049 | -0.563 | 0.007 | -0.591 | 0.004 | -0.660 | 0.002 | -0.546 | 0.005 |
|  |  | 5HT6 | 0.255 | 0.173 | 0.340 | 0.055 | 0.226 | 0.224 | 0.382 | 0.024 | 0.233 | 0.202 |
|  |  | 5HT7 | 0.359 | 0.069 | 0.434 | 0.017 | 0.418 | 0.016 | 0.564 | 0.002 | 0.370 | 0.038 |

Table S2. The summary of partial correlations between transcriptomic profiles and MRS neurometabolites controlling age and sex.

**Supplementary Table 3**

| Receptor Class | Subtype/Group |  | Semantic ACC | | Semantic RT | | Control ACC | | Control RT | |
| --- | --- | --- | --- | --- | --- | --- | --- | --- | --- | --- |
|  |  |  | r | qFDR | r | qFDR | r | qFDR | r | qFDR |
| GABAergic | GABA_A_ | GABRA1 | -0.477 | 0.028 | -0.046 | 0.902 | 0.070 | 0.852 | 0.124 | 0.917 |
|  |  | GABRA2 | 0.414 | 0.028 | -0.024 | 0.952 | -0.115 | 0.685 | -0.129 | 0.917 |
|  |  | GABRA3 | 0.383 | 0.045 | -0.015 | 0.952 | -0.124 | 0.685 | -0.127 | 0.917 |
|  |  | GABRA4 | -0.349 | 0.066 | 0.056 | 0.892 | 0.105 | 0.707 | 0.061 | 0.917 |
|  |  | GABRA5 | 0.370 | 0.057 | -0.041 | 0.908 | -0.142 | 0.680 | -0.096 | 0.917 |
|  |  | GABRB1 | 0.421 | 0.028 | 0.036 | 0.908 | -0.100 | 0.720 | -0.106 | 0.917 |
|  |  | GABRB2 | -0.474 | 0.028 | -0.040 | 0.908 | 0.052 | 0.879 | 0.095 | 0.917 |
|  |  | GABRB3 | 0.318 | 0.086 | -0.091 | 0.835 | -0.168 | 0.675 | -0.089 | 0.917 |
|  |  | GABRG1 | 0.360 | 0.059 | 0.140 | 0.762 | 0.026 | 0.917 | -0.132 | 0.917 |
|  |  | GABRG2 | -0.425 | 0.028 | -0.291 | 0.723 | -0.111 | 0.685 | -0.004 | 0.980 |
|  |  | GABRG3 | -0.454 | 0.028 | -0.055 | 0.892 | 0.065 | 0.854 | 0.117 | 0.917 |
|  | GABA_B_ | GABBR1 | 0.455 | 0.028 | 0.157 | 0.730 | -0.037 | 0.902 | -0.053 | 0.917 |
|  |  | GABBR2 | 0.390 | 0.043 | 0.134 | 0.783 | -0.046 | 0.879 | -0.025 | 0.923 |
| Glutamatergic | NMDA | GRIN1 | -0.063 | 0.741 | 0.185 | 0.723 | 0.166 | 0.675 | 0.104 | 0.917 |
|  |  | GRIN2A | -0.316 | 0.088 | 0.108 | 0.810 | 0.158 | 0.675 | 0.109 | 0.917 |
|  |  | GRIN2B | 0.297 | 0.105 | 0.100 | 0.835 | -0.072 | 0.852 | -0.038 | 0.917 |
|  |  | GRIN2C | 0.104 | 0.602 | 0.164 | 0.723 | 0.179 | 0.675 | 0.103 | 0.917 |
|  | AMPA | GRIA1 | 0.340 | 0.072 | -0.092 | 0.835 | -0.170 | 0.675 | -0.105 | 0.917 |
|  |  | GRIA2 | 0.346 | 0.069 | -0.080 | 0.870 | -0.174 | 0.675 | -0.105 | 0.917 |
|  |  | GRIA3 | 0.161 | 0.408 | -0.166 | 0.723 | -0.257 | 0.675 | -0.117 | 0.917 |
|  |  | GRIA4 | 0.457 | 0.028 | 0.247 | 0.723 | 0.168 | 0.675 | -0.008 | 0.971 |
|  | Kainate | GRIK1 | 0.297 | 0.105 | 0.228 | 0.723 | 0.045 | 0.879 | -0.050 | 0.917 |
|  |  | GRIK2 | 0.336 | 0.075 | -0.071 | 0.870 | -0.120 | 0.685 | -0.140 | 0.917 |
|  |  | GRIK3 | 0.255 | 0.161 | -0.006 | 0.978 | -0.151 | 0.675 | -0.104 | 0.917 |
|  |  | GRIK4 | 0.357 | 0.060 | -0.018 | 0.952 | -0.150 | 0.675 | -0.041 | 0.917 |
|  |  | GRIK5 | 0.436 | 0.028 | 0.131 | 0.783 | -0.033 | 0.908 | -0.088 | 0.917 |
|  | mGlu | GRM1 | 0.267 | 0.142 | 0.310 | 0.723 | 0.136 | 0.685 | 0.058 | 0.917 |
|  |  | GRM2 | -0.002 | 0.988 | -0.291 | 0.723 | -0.161 | 0.675 | -0.073 | 0.917 |
|  |  | GRM3 | 0.198 | 0.286 | 0.327 | 0.723 | 0.163 | 0.675 | 0.092 | 0.917 |
|  |  | GRM4 | -0.307 | 0.097 | -0.142 | 0.762 | -0.115 | 0.685 | 0.020 | 0.939 |
|  |  | GRM5 | 0.266 | 0.142 | -0.097 | 0.835 | -0.193 | 0.675 | -0.103 | 0.917 |
|  |  | GRM6 | 0.409 | 0.031 | 0.244 | 0.723 | 0.122 | 0.685 | 0.057 | 0.917 |
|  |  | GRM7 | -0.054 | 0.775 | -0.248 | 0.723 | -0.247 | 0.675 | -0.047 | 0.917 |
|  |  | GRM8 | -0.040 | 0.822 | -0.053 | 0.892 | -0.147 | 0.679 | -0.029 | 0.923 |
| Cholinergic | Nicotinic (nACh) | CHRNA2 | -0.285 | 0.121 | -0.108 | 0.810 | 0.009 | 0.969 | 0.155 | 0.917 |
|  |  | CHRNA3 | 0.329 | 0.077 | -0.109 | 0.810 | -0.186 | 0.675 | -0.131 | 0.917 |
|  |  | CHRNA4 | 0.231 | 0.209 | -0.158 | 0.730 | -0.246 | 0.675 | -0.147 | 0.917 |
|  |  | CHRNA5 | -0.325 | 0.079 | 0.049 | 0.902 | 0.152 | 0.675 | 0.111 | 0.917 |
|  |  | CHRNA6 | -0.091 | 0.636 | 0.191 | 0.723 | 0.248 | 0.675 | 0.158 | 0.917 |
|  |  | CHRNA7 | 0.269 | 0.140 | 0.299 | 0.723 | 0.208 | 0.675 | 0.025 | 0.923 |
|  |  | CHRNA9 | 0.237 | 0.197 | -0.146 | 0.762 | -0.228 | 0.675 | -0.147 | 0.917 |
|  |  | CHRNA10 | 0.246 | 0.178 | 0.177 | 0.723 | 0.021 | 0.933 | 0.034 | 0.917 |
|  |  | CHRNB2 | 0.224 | 0.222 | -0.168 | 0.723 | -0.239 | 0.675 | -0.117 | 0.917 |
|  |  | CHRNB3 | -0.104 | 0.602 | 0.115 | 0.810 | 0.236 | 0.675 | 0.035 | 0.917 |
|  | Muscarinic (mACh) | CHRM1 | -0.220 | 0.229 | 0.221 | 0.723 | 0.230 | 0.675 | 0.116 | 0.917 |
|  |  | CHRM2 | -0.333 | 0.075 | -0.070 | 0.870 | 0.113 | 0.685 | 0.057 | 0.917 |
|  |  | CHRM3 | 0.022 | 0.902 | 0.240 | 0.723 | 0.213 | 0.675 | 0.016 | 0.945 |
|  |  | CHRM4 | 0.324 | 0.079 | -0.121 | 0.810 | -0.178 | 0.675 | -0.113 | 0.917 |
|  |  | CHRM5 | -0.218 | 0.229 | -0.257 | 0.723 | -0.185 | 0.675 | -0.036 | 0.917 |
| Adrenergic | ɑ-Adrenergic | ADRA1D | -0.465 | 0.028 | -0.071 | 0.870 | 0.004 | 0.981 | 0.122 | 0.917 |
|  |  | ADRA1B | 0.282 | 0.122 | 0.343 | 0.723 | 0.253 | 0.675 | 0.080 | 0.917 |
|  |  | ADRA1A | 0.120 | 0.552 | -0.192 | 0.723 | -0.206 | 0.675 | -0.087 | 0.917 |
|  |  | ADRA2A | -0.421 | 0.028 | -0.060 | 0.892 | 0.092 | 0.754 | 0.117 | 0.917 |
|  |  | ADRA2B | 0.097 | 0.613 | -0.234 | 0.723 | -0.043 | 0.879 | -0.122 | 0.917 |
|  |  | ADRA2C | 0.418 | 0.028 | 0.017 | 0.952 | -0.062 | 0.857 | -0.140 | 0.917 |
|  | β-Adrenergic | ADRB1 | -0.368 | 0.058 | 0.054 | 0.892 | 0.167 | 0.675 | 0.114 | 0.917 |
|  |  | ADRB2 | 0.403 | 0.033 | -0.020 | 0.952 | -0.128 | 0.685 | -0.132 | 0.917 |
|  |  | ADRB3 | -0.099 | 0.613 | -0.226 | 0.723 | -0.264 | 0.675 | -0.052 | 0.917 |
| Dopaminergic | Dopamine | DRD1 | -0.300 | 0.105 | -0.181 | 0.723 | 0.030 | 0.910 | 0.062 | 0.917 |
|  |  | DRD2 | 0.142 | 0.463 | -0.165 | 0.723 | -0.245 | 0.675 | -0.116 | 0.917 |
|  |  | DRD3 | 0.051 | 0.781 | -0.253 | 0.723 | -0.249 | 0.675 | -0.124 | 0.917 |
|  |  | DRD4 | -0.069 | 0.733 | -0.119 | 0.810 | 0.048 | 0.879 | 0.077 | 0.917 |
| Serotonergic | 5-HT | 5HT1A | 0.440 | 0.028 | 0.149 | 0.762 | -0.045 | 0.879 | -0.046 | 0.917 |
|  |  | 5HT1B | 0.186 | 0.319 | -0.189 | 0.723 | -0.228 | 0.675 | -0.095 | 0.917 |
|  |  | 5HT1D | 0.416 | 0.028 | 0.171 | 0.723 | 0.065 | 0.854 | -0.045 | 0.917 |
|  |  | 5HT1E | -0.155 | 0.422 | 0.203 | 0.723 | 0.198 | 0.675 | 0.080 | 0.917 |
|  |  | 5HT1F | -0.434 | 0.028 | 0.004 | 0.978 | 0.113 | 0.685 | 0.119 | 0.917 |
|  |  | 5HT2A | -0.107 | 0.602 | 0.206 | 0.723 | 0.155 | 0.675 | 0.130 | 0.917 |
|  |  | 5HT2B | -0.064 | 0.741 | -0.186 | 0.723 | -0.009 | 0.969 | 0.046 | 0.917 |
|  |  | 5HT2C | 0.343 | 0.070 | -0.091 | 0.835 | -0.141 | 0.680 | -0.143 | 0.917 |
|  |  | 5HT3A | 0.364 | 0.059 | -0.072 | 0.870 | -0.133 | 0.685 | -0.093 | 0.917 |
|  |  | 5HT4 | 0.279 | 0.126 | -0.074 | 0.870 | -0.202 | 0.675 | -0.087 | 0.917 |
|  |  | 5HT5A | -0.434 | 0.028 | -0.037 | 0.908 | 0.112 | 0.685 | 0.121 | 0.917 |
|  |  | 5HT6 | 0.151 | 0.430 | -0.214 | 0.723 | -0.234 | 0.675 | -0.112 | 0.917 |
|  |  | 5HT7 | 0.289 | 0.115 | -0.108 | 0.810 | -0.191 | 0.675 | -0.097 | 0.917 |

Table S3. The summary of partial correlations between transcriptomic profiles and task performance controlling age and sex.

**Supplementary Table 4**

| Mediator | Receptor Class | Subtype | Path a Effect | Path a p | Path b Effect | Path b p | Total effect (c) | Total Effect p | Direct Effect (c') | Direct effect p | Indirect effect (a x b) | 95% Bootstrap CI | | Proportion Mediated (PM) |
| --- | --- | --- | --- | --- | --- | --- | --- | --- | --- | --- | --- | --- | --- | --- |
| GABA+ | GABAergic | GABRA1 | -6.023 | 0.007 | 6.801 | 0.033 | -103.458 | 0.012 | -62.495 | 0.139 | -40.964 | [-96.6430 | -6.4510] | 40% |
|  |  | GABRA2 | 4.138 | 0.012 | 7.799 | 0.017 | 57.575 | 0.062 | 25.302 | 0.412 | 32.273 | [6.3360 | 84.0013] | 56% |
|  |  | GABRA3 | 5.909 | 0.023 | 7.850 | 0.014 | 87.675 | 0.070 | 41.290 | 0.384 | 46.385 | [8.2659 | 133.8723] | 53% |
|  |  | GABRA4 | -7.829 | 0.050 | 8.327 | 0.009 | -103.510 | 0.163 | -38.326 | 0.588 | -65.185 | [-185.3204 | -2.8309] | 63% |
|  |  | GABRA5 | 7.080 | 0.022 | 8.325 | 0.010 | 84.834 | 0.143 | 25.892 | 0.647 | 58.942 | [10.6958 | 166.7172] | 69% |
|  |  | GABRB1 | 5.317 | 0.027 | 7.516 | 0.017 | 91.345 | 0.038 | 51.385 | 0.234 | 39.960 | [6.6849 | 107.7987] | 44% |
|  |  | GABRB2 | -5.183 | 0.010 | 6.994 | 0.028 | -89.008 | 0.016 | -52.760 | 0.159 | -36.247 | [-89.1621 | -6.4862] | 41% |
|  |  | GABRB3 | 4.808 | 0.033 | 8.738 | 0.007 | 46.875 | 0.267 | 4.866 | 0.905 | 42.009 | [7.3710 | 127.3897] | 90% |
|  |  | GABRG3 | -3.893 | 0.011 | 7.069 | 0.026 | -65.943 | 0.019 | -38.422 | 0.175 | -27.521 | [-64.7408 | -5.0888] | 42% |
|  |  | GABBR1 | 3.635 | 0.029 | 6.939 | 0.023 | 76.420 | 0.011 | 51.196 | 0.082 | 25.224 | [3.9731 | 64.4377] | 33% |
| Glx | GABAergic | GABRA1 | -26.868 | 0.002 | 1.354 | 0.079 | -120.406 | 0.003 | -84.036 | 0.048 | -36.370 | [-86.3882 | -1.2581] | 30% |
|  |  | GABRA2 | 18.650 | 0.003 | 1.599 | 0.043 | 72.004 | 0.014 | 42.174 | 0.170 | 29.830 | [3.9313 | 72.6556] | 41% |
|  |  | GABRA3 | 26.038 | 0.010 | 1.689 | 0.029 | 106.045 | 0.023 | 62.069 | 0.188 | 43.976 | [6.8286 | 110.5481] | 41% |
|  |  | GABRA4 | -39.915 | 0.014 | 1.706 | 0.026 | -166.095 | 0.025 | -97.987 | 0.188 | -68.108 | [-186.4709 | -7.9082] | 41% |
|  |  | GABRA5 | 26.002 | 0.027 | 1.747 | 0.021 | 115.829 | 0.031 | 70.402 | 0.182 | 45.427 | [5.6247 | 111.7942] | 39% |
|  |  | GABRB1 | 28.668 | 0.002 | 1.570 | 0.051 | 106.747 | 0.012 | 61.735 | 0.175 | 45.012 | [4.4277 | 112.4154] | 42% |
|  |  | GABRB2 | -23.627 | 0.003 | 1.376 | 0.072 | -107.581 | 0.003 | -75.070 | 0.048 | -32.511 | [-78.7403 | -1.2533] | 30% |
|  |  | GABRB3 | 19.036 | 0.027 | 1.866 | 0.016 | 69.607 | 0.078 | 34.086 | 0.380 | 35.521 | [4.9402 | 87.4833] | 51% |
|  |  | GABRG1 | 27.678 | 0.026 | 1.760 | 0.020 | 120.128 | 0.034 | 71.414 | 0.202 | 48.714 | [7.3332 | 114.9872] | 41% |
|  |  | GABRG2 | -42.470 | 0.010 | 1.533 | 0.042 | -203.118 | 0.006 | -138.033 | 0.068 | -65.086 | [-153.2057 | -6.5115] | 32% |
|  |  | GABRG3 | -19.296 | 0.001 | 1.437 | 0.072 | -76.105 | 0.006 | -48.380 | 0.104 | -27.726 | [-66.7796 | -1.7213] | 36% |
|  |  | GABBR1 | 19.178 | 0.003 | 1.506 | 0.055 | 78.748 | 0.007 | 49.872 | 0.109 | 28.876 | [0.1986 | 68.8326] | 37% |
|  |  | GABBR2 | 17.797 | 0.011 | 1.718 | 0.027 | 71.050 | 0.028 | 40.482 | 0.214 | 30.568 | [4.1628 | 72.6289] | 43% |
|  | Glutamatergic | GRIN2A | -22.360 | 0.011 | 1.833 | 0.020 | -76.535 | 0.061 | -35.559 | 0.387 | -40.976 | [-123.6192 | -4.3315] | 54% |
|  |  | GRIA1 | 12.152 | 0.012 | 1.821 | 0.020 | 42.664 | 0.057 | 20.530 | 0.362 | 22.134 | [3.6351 | 56.2082] | 52% |
|  |  | GRIA2 | 18.097 | 0.006 | 1.828 | 0.022 | 58.623 | 0.057 | 25.546 | 0.419 | 33.077 | [4.8682 | 79.2039] | 56% |
|  |  | GRIA4 | 34.928 | 0.003 | 1.447 | 0.063 | 149.378 | 0.005 | 98.831 | 0.079 | 50.546 | [0.7903 | 116.1219] | 34% |
|  |  | GRIK1 | 51.446 | 0.012 | 1.908 | 0.016 | 157.821 | 0.097 | 59.656 | 0.531 | 98.165 | [14.0398 | 232.7442] | 62% |
|  |  | GRIK2 | 23.451 | 0.030 | 1.805 | 0.018 | 97.325 | 0.049 | 55.000 | 0.256 | 42.325 | [5.3148 | 115.3897] | 43% |
|  |  | GRIK4 | 23.016 | 0.025 | 1.805 | 0.018 | 92.523 | 0.050 | 50.976 | 0.273 | 41.547 | [6.7698 | 99.7378] | 45% |
|  |  | GRIK5 | 20.284 | 0.002 | 1.521 | 0.057 | 77.915 | 0.009 | 47.057 | 0.142 | 30.858 | [0.3400 | 74.0776] | 40% |
|  |  | *GRM6* | 45.611 | 0.007 | 1.623 | 0.036 | 189.859 | 0.015 | 115.836 | 0.146 | 74.024 | [9.8864 | 169.2652] | 39% |

Table S4. Mediation analyses of GABA+ and glutamate–glutamine Glx linking neurotransmitter receptor-subunit gene expression to semantic accuracy.
